# Profiling Expression Strategies for a Type III Polyketide Synthase in a Lysate-Based, Cell-free System

**DOI:** 10.1101/2023.11.30.569483

**Authors:** Tien T. Sword, Jaime Lorenzo N. Dinglasan, Ghaeath S.K. Abbas, J. William Barker, Madeline E. Spradley, Elijah R. Greene, Damian S. Gooden, Scott J. Emrich, Michael A. Gilchrist, Mitchel J. Doktycz, Constance B. Bailey

## Abstract

Some of the most metabolically diverse species of bacteria (e.g., Actinobacteria) have higher GC content in their DNA, differ substantially in codon usage, and have distinct protein folding environments compared to tractable expression hosts like *Escherichia coli*. Consequentially, expressing biosynthetic gene clusters (BGCs) from these bacteria in *E. coli* frequently results in a myriad of unpredictable issues with protein expression and folding, delaying the biochemical characterization of new natural products. Current strategies to achieve soluble, active expression of these enzymes in tractable hosts, such as BGC refactoring, can be a lengthy trial-and-error process. Cell-free expression (CFE) has emerged as 1) a valuable expression platform for enzymes that are challenging to synthesize *in vivo*, and as 2) a testbed for rapid prototyping that can improve cellular expression. Here, we use a type III polyketide synthase from *Streptomyces griseus*, RppA, which catalyzes the formation of the red pigment flaviolin, as a reporter to investigate BGC refactoring techniques. We synergistically tune promoter and codon usage to improve flaviolin production from cell-free expressed RppA. We then assess the utility of cell-free systems for prototyping these refactoring tactics prior to their implementation in cells. Overall, codon harmonization improves natural product synthesis more than traditional codon optimization across cell-free and cellular environments. Refactoring promoters and/or coding sequences via CFE can be a valuable strategy to rapidly screen for catalytically functional production of enzymes from BCGs. By showing the coordinators between CFE versus *in vivo* expression, this work advances CFE as a tool for natural product research.

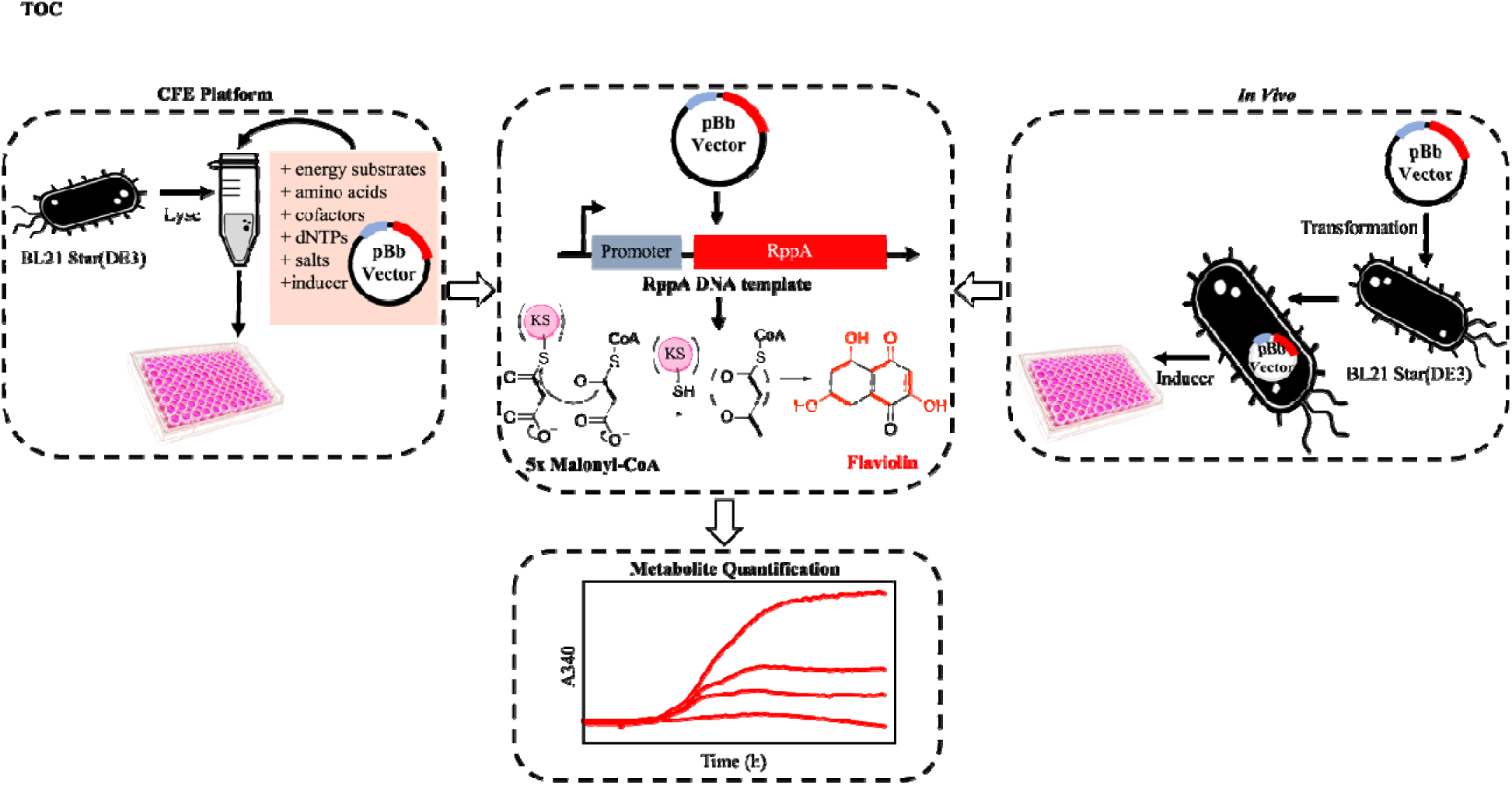

## Introduction

Microbial secondary metabolism generates a vast number of complex secondary metabolites, or natural products, with varied chemistry. ^1^ Members of the order Actinomycetales, and especially those from the genus *Streptomyces,* have been shown to be a valuable lineage in terms of their capacity for secondary metabolism. ^2–5^ Indeed, *Streptomyces* genomes are known for harboring a large number of biosynthetic gene clusters (BGCs) that are attractive for the discovery of novel enzymes and their chemical products. Unfortunately, these positive aspects of *Streptomyces spp.* are offset by their slow doubling time, mycelial clumping, thick cell walls, high GC content (∼70-75%), and relatively cumbersome genetics. Consequently, biochemists who study *Streptomyces spp.-*derived natural products struggle to generate enough protein for biochemical or structural characterization. As a result, most efforts for expressing proteins of interest as soluble, functional constructs are done in tractable hosts like *E. coli*, rather than *Streptomyces spp*. Because of the deep evolutionary divergence between Actinobacteria and Proteobacteria such as *E. coli*, the differences in their metabolic backgrounds, dissimilar codon usage and genome attributes, as well as protein folding environments, expression of heterologous genes from *Streptomyces spp.* and, in turn, product synthesis often fails. ^6^ Where BGC expression in *E. coli* is possible, a myriad of parameters usually require optimization. Oftentimes, using the native coding sequence amplified from genome DNA with common expression systems (e.g., the pET expression system) ^7,8,9^ results in poor expression or inclusion body formation. ^10^ In other cases, product toxicity can impede the use of *E. coli* strains without engineering host tolerance. Finding suitable recombinant expression systems therefore involves screening of refactoring choices that include choice of regulatory elements like promoters, and of coding strategy. This process can be time-consuming and laborious, a significant bottleneck to researcher workflows.

Cell-free expression (CFE) platforms employ either crude cell lysates (or extracts) or an *in vitro* TX-TL PURE system and can bypass limitations in secondary metabolite production observed in in vivo expression systems.^11,12^ The TX-TL PURE system or PURExpress employ the minimal number of recombinant elements required for transcription and translation (TX-TL) while approaches using crude cell lysates are derived from intact living cells. Harnessing the TX-TL machinery preserved in lysates allows protein expression in the absence of other normal cellular functions, a feature that can be leveraged to manufacture enzymes that are difficult to synthesize in microbial hosts. Lysates also retain metabolic pathways that can be engineered to accumulate precursor molecules for heterologous biosynthetic enzymes.^13–15^ CFE systems are thus often used for synthesizing BGCs and their product metabolites, especially when cytotoxicity represents a limiting factor to heterologous *in vivo* production.^16–18^ Additionally, CFE systems can be used to optimize the cell-based expression of soluble BGCs when leveraged as testbeds for genetic refactorization. Prototyping different genetic constructs in these platforms is relatively rapid as it bypasses time limitations associated with culturing and genetically manipulating live cells. To these ends, investigating refactoring strategies in a cell-free environment can benefit development of cell-free and cell-based BGC expression platforms.

While optimizations of protein expression via refactoring have usually focused on robustly expressed reporters (e.g., sfGFP), ^19–24^ we sought to evaluate common refactoring parameters using a reporter that was relevant to the enzymatic activity of genes involved in to secondary metabolite formation. ^7^ To further explore the ability to refactor for functional catalytic activity, we chose a model protein, RppA (40.1 kDa), which is a type III polyketide synthase from *Streptomyces griseus* that generates flaviolin, which is a red pigment that has limited catalytically functional expression in *E. coli* even with current optimization. ^25–27^ Using flaviolin production as a reporter for catalytically functional RppA expression, we varied parameters that are relevant to improving expression. This included the use of three different, commonly used inducible promoters: the workhorse T7/lac system, the pBAD arabinose promoter, ^28^ and the pTet anhydrotetracycline promoter. ^29^ In addition to varying promoters, we also evaluated the impact of four different methods for designing synonymous coding sequences for generating higher levels of heterologous gene expression by using codons that better reflect the host’s (*E. coli*’s) codon preferences. Notably, while used extensively *in vivo,* inducible promoters beyond the T7 system have not been extensively investigated in CFE. We also demonstrate strong positive correlations between cell-free and cell-based expression and discuss the feasibility of this approach for refactoring challenging proteins *in vitro* prior to *in vivo* production. Taken together, this work demonstrates a coordinated strategy to apply a lysate-based cell-free environment for profiling genetic constructs that promote catalytically active enzyme formation. The results are applicable to both cell-free and cell-based systems and can be used to generate biosynthetic proteins for characterizing elements of engineered biosynthetic pathways.

## Results and discussion

### Design of the vector system

To design a library of constructs to improve expression, we focused on two commonly varied refactoring parameters: promoter choice and codon usage. We used four distinct coding sequences and four distinct promoters (**Figure 1A**). In terms of promoter choice, while the T7-lac promoter ^30^ combination is the basis of the most commonly used expression system in *E. coli* (particularly the pET expression system) ^8,9^ other promoters that are not strong as the T7 promoter nor as leaky as the ITPG inducible lac operon are sometimes used to promote soluble expression of challenging to express proteins *in vivo*. ^30^ Other commonly used vectors include the pTet promoter, which is anhydrotetracycline inducible, ^30^ and the pBAD promoter that is arabinose inducible ^28^ both of which are less leaky than the lac operon (**Figure 1B**). To obtain these constructs, we used a series of BioBrick vectors designed by Keasling and coworkers ^30^ that included vector pBbE2k (harboring the pTet promoter), and pBbE8k (harboring the pBAD promoter), and pBbE7k (harboring the T7 promoter under control of the lac operon).

**Figure 1.**
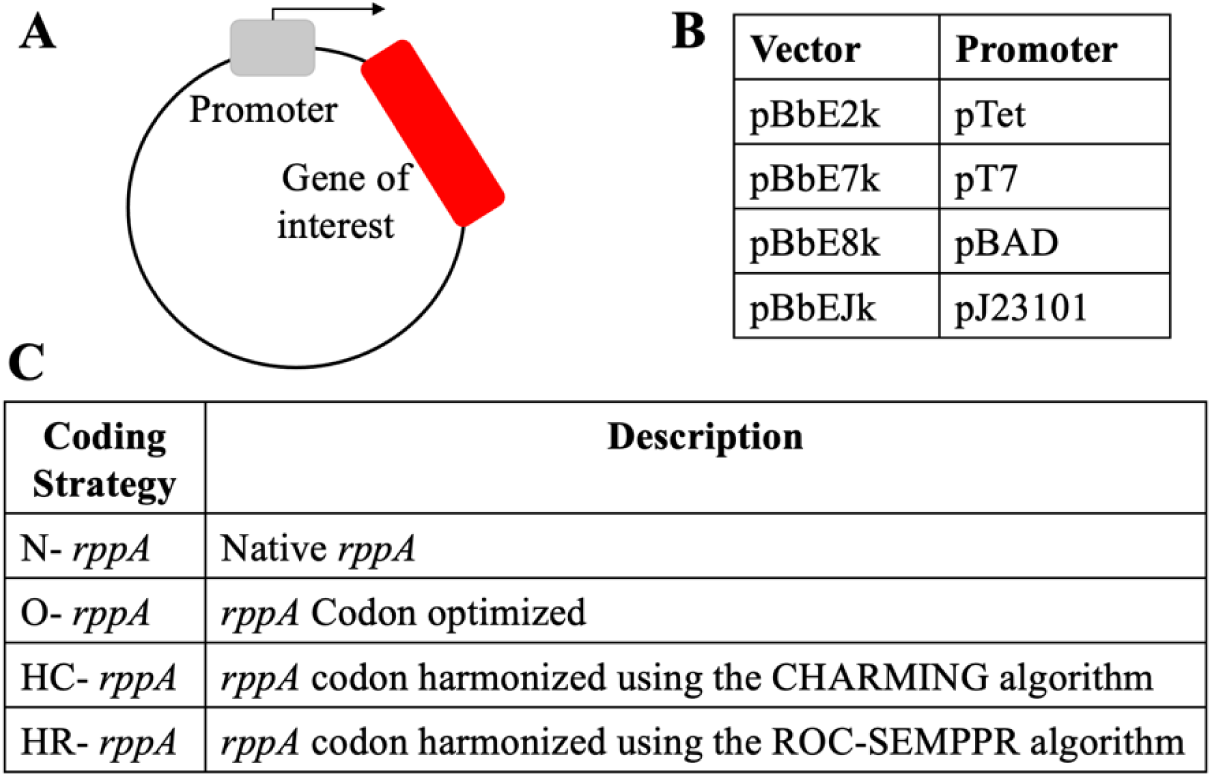
Design and nomenclature of plasmids in this study. (A) Each plasmid tested is composed of two modules: a backbone containing different promoters and the gene of interest (*rppA*) (B) Names of the BioBrick vectors (pBb) carrying different promoters. ^30^ (C) Names of the varied coding sequences.

To complement this series of promoters, we varied synonymous coding sequences. When expressing proteins from a high GC bacterium that differs substantially from *E. coli* in terms of codon usage, there are several approaches that can be taken. While sometimes the natural coding sequence results in successful protein expression in *E. coli*, protein expression and folding issues can occur. These issues with expression and poor folding/solubility can originate from a mismatch between tRNA pools typical of each organism, which can change the rate of translation (e.g., cause stalls at the ribosome) and potentially disrupt appropriate co-translational folding. ^10,31–33^ Codon optimization is a common strategy to alter codon assignments, and appropriate algorithms are readily accessible from most commercial gene synthesis companies via replacement by codons used more frequently in the host’s genome or transcriptome.^34–40^ While these genome and transcriptome optimized sequences can aid in successful expression of the heterologous product, improperly folded products can still result.^41,42^ Another approach is codon “harmonization,” which has been posited to improve expression via better co-translational folding. ^43^ Codon harmonization involves identifying and replicating patterns of codon usage in the donor organism with comparable patterns of codon usage in the heterologous host. ^44,4544^ Typically, synonymous codons (or sliding windows of these codons) are assigned computational estimates of their frequency of appearance in the original host organism. Next, single codons are changed based on frequencies estimated in the desired heterologous host to better replicate the source organism’s frequency patterns, which enables stalling patterns at the ribosome that are more akin to how they originally evolved and therefore might result in proper protein folding.^45^

To explore the effect of synonymous coding, four constructs were compared. The native coding sequence amplified directly from *Streptomyces griseus* genomic DNA, routine codon optimization was performed by Integrated DNA Technology’s (IDT) codon optimization algorithm, and finally, two codon harmonization constructs were designed, the first using the CHARMING (HC-*rppA*) and a new method based on ROC-SEMPPR (HR-*rppA*) (**Figure 1C**). Briefly, the CHARMING method applies a relative measure of codon usage called “% MinMax” (%MM). ^42^ %MM values are computed as described by Chaney et al.^33,46,47^ using overall codon usage from an organism obtainable from various sources including codon usage information tabulated in the international DNA sequence database (Kazusa: https://www.kazusa.or.jp/codon/).^48^ CHARMING uses a sliding window to estimate %MM-based deviations between the original and target organism codon usage for a given protein. While large deviations exist within one or more windows, single synonymous changes are made that best “harmonize” the values, i.e., reduce the overall %MM difference in that specific window. In short, this algorithm will minimize the sum of | MM_original – MM_target | over all windows and will proceed until five consecutive iterations where no beneficial, i.e., reduces differences between the %MM values, changes are found. The size of the windows used was the CHARMING default value, which was set to be most consistent with ribosome fingerprint-based pausing estimates.^42^ In contrast, ROC-SEMPPR takes an evolutionary approach to estimating the translational efficiency of an amino acid’s synonymous codons within a given organism. ROC-SEMPPR does so by fitting a probabilistic, population genetics based model of sequence evolution, which includes the contributions of selection, mutation bias, and genetic drift, to an organism’s coding sequences. ^49,50^ By simultaneously analyzing intragenic and intergenic patterns of synonymous codon usage within a genome, ROC-SEMPPR estimates differences in ribosome pausing times among synonymous codons translational efficiency and mutation bias between codons and differences in protein production rates between genes. Fitting ROC-SEMPPR separately to the donor (in this case *Streptomyces griseus*) and host genomes (in thi case *E. coli)* enables the ability to rank each amino acid’s synonymous codons by their translational efficiencies within the donor and host, respectively.

The promoter and coding sequence combinations represent a total of 16 different constructs. The naming convention for the components of the 16 constructs are described in **Figure 1**.

### Initial cell-free experiments for a type III PKS enable the production of flaviolin

RppA catalyzes polyketide synthesis by condensing and cyclizing five molecules of the pentaketide polyketide tetrahydroxynapthalene (THN). ^25,26^ Subsequently, THN undergoes a spontaneous oxidation reaction to convert THN to flaviolin (**Figure 2A**). The formation of th red-brown flaviolin pigment can thus be monitored as it readily absorbs light at 340 nm. ^27^ As metabolite production in an *E. coli* lysate-based cell-free system is correlated with the amount of protein that is expressed, ^51^ we used the amount of flaviolin produced as a proxy for estimating catalytically functional RppA production. To establish assay conditions, we first sought to determine the threshold for pigment production against a cell lysate background. To do so, we spiked purified flaviolin into lysate preparations in the absence of DNA. Sufficient pigment concentrations could be detected at the micromolar range, demonstrating sufficient sensitivity to proceed (**Figure S1**). With an assay established, pigment production could be tested at variable temperature conditions and plasmid DNA concentrations.

**Figure 2.**
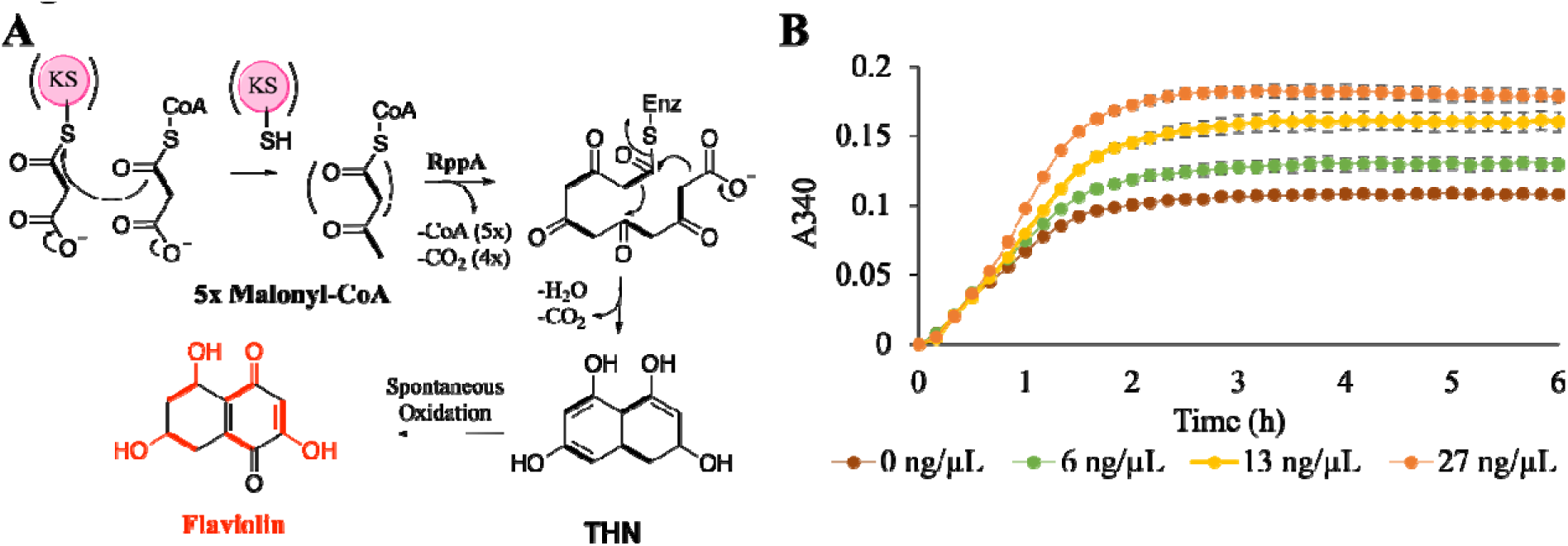
Flaviolin production via RppA in CFE reactions. (A) Schematic of flaviolin biosynthetic pathway by type III PKS, RppA. (B) CFE reactions were initiated with increasing concentrations of codon optimized *rppA* (o-*rppA*) driven by pT7 promoter in plasmid DNA, resulting in increasing flaviolin production (observed by increase of absorbance at 340 nm).

To define temperature conditions for conducting CFE experiments, we used the pET28b expression plasmid and codon-optimized *rppA* (O-*rppA*) in a lysate-based system using *E. coli* BL21 Star(DE3), a BL21(DE3) variant that has the DE3 lysogen under control of a lacUV5 promoter. ^52^ We have previously had success using this lysate for the production of pigments from *Streptomyces.* ^51^ Our rationale for this initial choice of coding sequence and promoter was twofold: 1) prior studies from our laboratory suggest that heterologous expression of enzymes originating from *Streptomyces* that form pigments show improved expression and solubility when using *E. coli* optimized sequences ^10^ and 2) the T7 promoters (and specifically pET vectors) have been widely used in CFE. ^51,53–57^ These initial experiments revealed superior pigment production at 30°C, so we proceeded with 30°C for all subsequent experiments (**Figure S2**). To remove confounding variables from differing intergenic regions and a different origin of replication in the pET vector as compared to the BioBrick vectors, we repeated this experiment using pBbE7k as the backbone for consistency with the pBAD and pTet constructs. Increasing the amount of the plasmid construct pBbE7k-O-rppA (containing *E. coli* codon-optimized *rppA* driven by pT7 promoter) with *E. coli* BL21 Star(DE3) lysate containing endogenous IPTG at 30°C demonstrated that the DNA template concentration affects protein expression and therefore product formation (**Figure 2B**).

### Establishing the application of non-IPTG inducers for cell-free with other promoters

The ITPG inducible T7-lac expression system, especially the pET expression system is by far the most heavily used expression system for heterologous recombinant protein production in *E. coli* due to its engineered promoter strength. ^58^ However, its extreme promoter strength, combined with leakiness of the lac operon, adds to the metabolic burden of the cell resulting in decreased fitness less than optimal protein expression. For proteins that seem to be better expressed and appropriately folded with tighter transcriptional control, other inducible promoter systems have been developed, such as the arabinose inducible pBAD and anhydrotetracycline inducible pTet systems. Indeed, there is precedence for alternate promoter systems improving expression of challenging, complex proteins from *Streptomyces*; proteins from the borrelidin polyketide synthase from *Streptomyces parvulus* were found to have improved expression using the pTet promoter when compared to the T7 system. ^56,59,60^ Extensive efforts have been made to apply the T7 promoter series under the lac operator in CFE. ^53,61^ However, other inducible systems remain extremely underexplored, with only a few reports of their usage in lysate based CFE systems.^62–64^

To compare the performances of each promoter in a lysate-based system, extracts were first prepared from *E. coli* BL21 Star(DE3) in the absence of IPTG. While IPTG is typically added to BL21 Star(DE3) cultures to promote the expression of T7 RNA polymerase prior to cell lysis, we omitted this step to prepare a lysate background that is appropriate for the comparison of non-IPTG inducible promoters. Using this batch of lysate, reactions were then first optimized to express RppA under pT7 and with the supplementation of IPTG to the lysate reaction. IPTG concentrations were supplemented to reactions containing 50 ng/µL T7 RNA polymerase. ^65^ Maximal flaviolin production was observed with a concentration of 500 µM IPTG (**Figure S3A, C**). The same lysate preparations were then used for protein expression driven by the pTet and pBAD promoters under different ranges of anhydrotetracycline and L-arabinose, respectively. However, under these conditions, we did not observe detectable flaviolin production. We hypothesized that flaviolin signals are not detectable under these conditions due to lower soluble protein expression, and, consequently, less flux being driven from endogenous malonyl-CoA precursor pools to flaviolin formation. Thus, we hypothesized that increasing substate availability would increase flaviolin production to detectable levels. To test this, malonyl-CoA was added to reactions containing pBbE7k-O-rppA plasmid DNA. Indeed, adding up to 500 µM malonyl-CoA to the reactions enabled higher levels of flaviolin synthesis (**Figure S3B, C**). *E. coli* can possess low levels of malonyl-CoA endogenously,^66^ therefore, with the pT7 system, there is a high level of RppA protein that flaviolin can be detectably produced even without malonyl-CoA supplementation. The low expression from the pTet/pBAD constructs combined with the low endogenous malonyl-CoA pools may be why the signals from pTet/pBAD constructs are low within the timeframe we measured. Applying this concentration (500 µM) of malonyl-CoA to systems driven by pTet and pBAD promoters results in the successful detection of flaviolin, albeit at later time points, confirming that soluble RppA expression is attainable with these promoters. A wide range of anhydrotetracycline and L-arabinose concentrations were tested using pBbE2k-O-*rppA* (O-*rppA* driven by pTet) and pBbE8k-O-*rppA* (O-*rppA* driven by pBAD), respectively (**Figures S4, S5**). For the pTet promoter, we observed the greatest expression level at 50 µM anhydrotetracycline (**Figure S4A, B**, whereas 10 mM L-arabinose was greatest for the pBAD promoter (**Figure S5A, B**.

Intriguingly, when optimized inducer concentrations are applied, overall flaviolin production is comparable between the pTet and pT7 conditions, even though synthesis is clearly delayed in the former system. Faster expression in the pT7 system could be due to promoter strength or the availability of exogenously supplied T7 RNA polymerase, whereas the other promoters rely on low levels of endogenous *E. coli* RNA polymerase. To confirm that fast expression from pT7 is a result of this promoter’s strength, we first supplied reactions with decreasing concentrations of T7 RNA polymerase. While overall flaviolin production decreases with lower polymerase levels, flaviolin synthesis still begins within the first hour under all conditions. These data imply that fast expression under pT7 is due to this system being less tightly regulated compared to pTet and pBAD and not necessarily the availability of the polymerase (**Figure S6**). To further interrogate this phenomenon, we tested our promoter strategy with sfGFP, allowing us to distinguish protein expression from enzyme catalysis or precursor availability. As an initial step, we verified that inducer concentrations performed best for RppA expression were also performed best for inducing sfGFP expression under the control of different promoters in CFE reactions (**Figure S7A-C**). We subsequently compared sfGFP synthesis from these inducible promoters to one another and to production from a constitutive promoter, pJ23101, in an analogous biobrick vector (pBbEJk). Like expression from pTet and pBAD, constitutive expression would also rely on endogenous *E. coli* RNA polymerase pools but should be quicker in theory because of a lack of regulation. Evidently, there is a delay in the expression of sfGFP under the control of either pTet, pBAD, and pJ23101 promoters compared to pT7 (**Figure S7D**). While the constitutive promoter produced relatively low levels of sfGFP, it did result sfGFP expression over a faster timeframe compared to pTet and pBAD (**Figure S7D & E**). Thus, delayed CFE from pTet and pBAD is likely a result of their tighter regulation compared to pT7.

### Synergistic Effect of Promoter and Coding Strategy in Refactoring Proteins for RppA CFE

After establishing a set of experimental conditions for detectable flaviolin formation driven by non-T7 inducible promoters, we sought to determine the synergistic effect of promoter plus coding strategy in refactoring a protein for optimized expression in our cell-free system. Overall, we found the strongest promoter-coding strategy to be the pBAD promoter using the ROC-SEMPPR method (HR-*rppA*) which is slightly higher than the CHARMING method (HC-*rppA*) driven by the pTet promoter and has significantly higher expression than all other constructs (**Figure 3A-C)**. Whether or not ROC-SEMPPR will perform similarly well in other situations remains to be determined. It does, however, suggest that naively ranking codons based on their occurrence in a genome (which ignores the role of mutation bias, variation in expression between genes) or transcriptome (which ignores the role of mutation bias and the limits drift places on adaptation), while useful, can be improved upon. In addition, we observed that expression with the pT7 promoter can be drastically improved when the lysate is supplied with endogenous IPTG as opposed to exogenous IPTG during production of the cell lysate to express the T7 RNAP in the BL21(DE3)Star strain (**Figure S6**). Interestingly, with the pT7 promoter, the HC construct produced the least amount of flaviolin, even lower than the natively coded sequence cloned from genomic DNA. The CHARMING method used to generate the HC construct uses sliding windows to estimate local rates of translation across the coding sequence, e.g., to best facilitate natural ribosomal stalling (see Methods); ^42^ however, because few rare codons are observed in the native *rppA* coding sequence, it appears that this method is not necessary to promote functional protein production using a pT7 promoter. (**Figure 3A**). The trend of each coding sequence considered is different for each promoter. For the pTet promoter, HC-*rppA* has the best expression, then HR-*rppA* is the second best, while the native construct (N-*rppA*) does not express at all (**Figure 3B**). Similarly, with the pBAD promoter, the native construct has low expression while HR-*rppA* has the best expression followed by HC*-rppA* (**Figure 3C**). Because these codon harmonization models do not account for inducible expression, we sought to determine the effects of these re-coding strategies on flaviolin production when RppA is expressed under pJ23101, allowing us to decouple expression from induction. In this case, HR-*rppA* expressed drastically better than other constructs, whereas, in contrast, the O-*rppA* did not express at all. Importantly, these experiments show that the choice of codon optimization/harmonization techniques and promoter both impact protein expression synergistically, which is an important consideration for future efforts to use CFE for high-yield protein expression. Additionally, unlike RppA, sfGFP expressed best under pT7 control while the constitutive promoter does not express well in these CFE conditions (**Figure S7D**). Thus, these results also confirm that different proteins have varying optimal CFE expression conditions. ^51^

**Figure 3.**
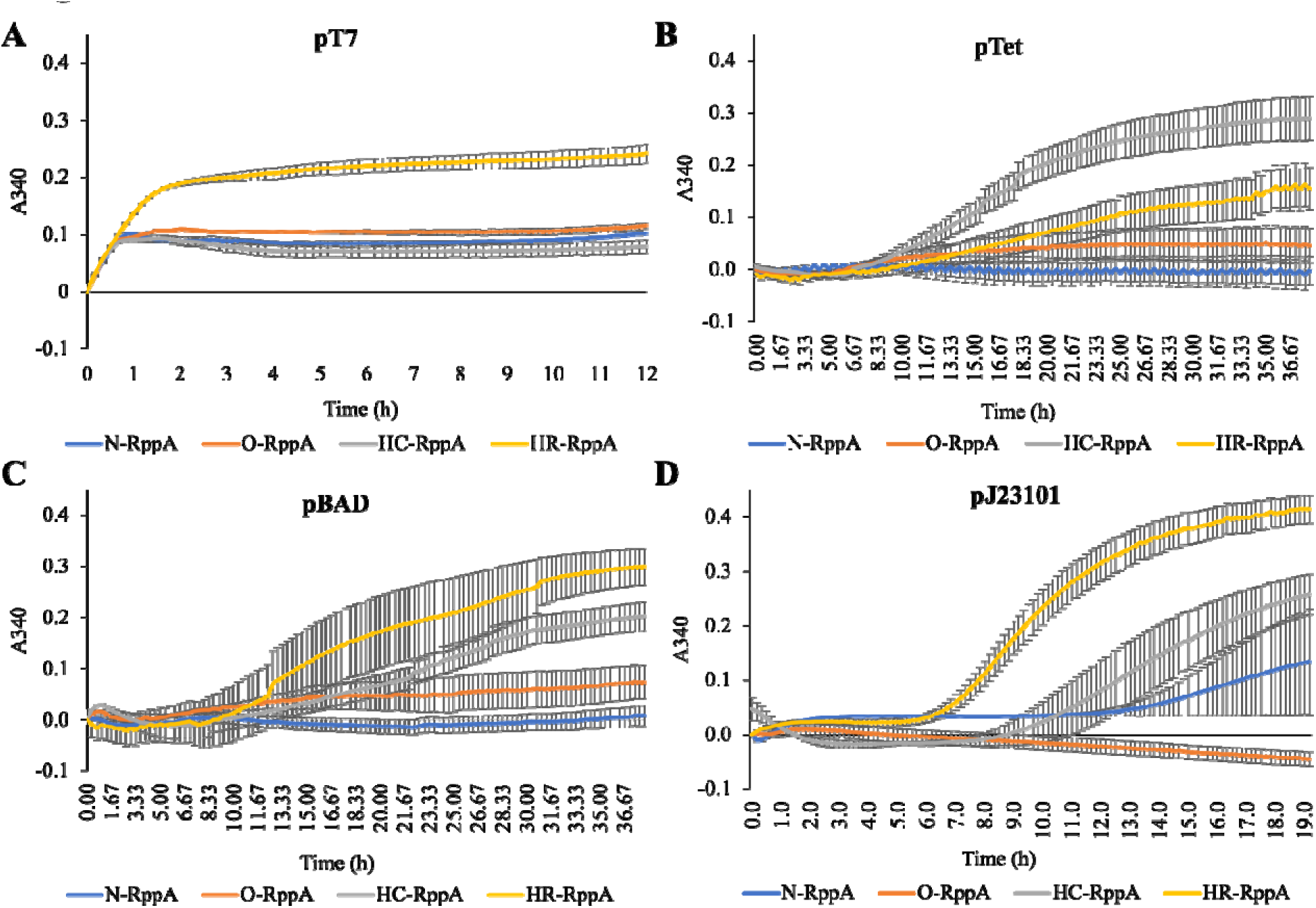
Flaviolin measurement in CFE reactions initiated with plasmids carrying different promoter and coding sequence combinations. Error bars represent the standard error of the mean (*n* = 3). (A) CFE reactions containing one of four pT7 promoter constructs, 50 ng/µL T7 RNA polymerase, and 500 µM IPTG. Reactions were run in triplicate and read every 10 mins for 12 hr. (B) CFE reactions containing one of four pTet promoter constructs, 500 µM malonyl-CoA, and 50 µM tetracycline. Reactions were run in triplicate and read every 20 mins for 38 hours. (C) CFE reaction containing four pBAD promoter constructs plasmid DNA, 500 µM malonyl-CoA, and 10 mM L-arabinose. Reactions were run in triplicate and read every 20 mins for 38 hours. (D) CFE reaction containing four pJ23101 promoter constructs plasmid DNA and 500 µM malonyl-CoA. Reactions were run in triplicate and read every 10 mins for 20 hours. All reactions were prepared with *E. coli* BL21 Star(DE3) lysates. Error bars represent the standard error of the mean (*n* = 3).

### Investigating the Utility of CFE for Prototyping Refactoring Techniques for *In vivo* Production

Expanding strategies for profiling expression choices to less explored choices (e.g., non-T7 promoters and lesser used refactoring strategies) and demonstrating their synergistic effects in CFE is more valuable when there are correlations between CFE and *in vivo*. ^54,67,68^ To determine whether the refactoring strategies we explored are correlative to *in vivo* expression, we transformed each of our refactored constructs into BL21Star (DE3) cells. First, OD_600_ of the codon-optimized construct of each promoter was measured to determine inducing time. Optimal OD_600_ for each promoter/operator combination was based on literature precedent for standard ODs of induction for each promoter respectively (OD_600_ = ∼0.8 for T7-lac, OD_600_ = ∼0.2 for pBAD, OD_600_ = ∼0.6 for pTet). ^69–71^ In a 96-well plate, from the initial culture with OD_600_ = 0.05, pT7 constructs take 3.5 hr to reach OD_600_ = 0.8, pTet constructs take 3 hours to reach OD_600_ = 0.6, and pBAD constructs take 70 mins to reach OD_600_ = 0.2. Next, we varied the inducer concentrations for each promoter – *rppA* DNA construct (**Figure S8**). Of the conditions we evaluated, we found that inducer concentrations are different with CFE reactions: IPTG concentration reaches greatest expression at 500 µM, anhydrotetracycline at 1000 nM, and L-arabinose at 5 mM. These conditions correlate with previous reported for RFP, thus, they were used for sfGFP expression. ^30^

When comparing the effect of codon optimization/harmonization on RppA expression, without considering promoter choice, trends only correlate between the *in vivo* and *in vitro* experiments in the pT7 and the constitutive promoter data (**Figures 3A&D, 4A&D**). When expressing with pTet, the HC-*rppA* outperforms HR-*rppA in vitro* while these two harmonized sequences perform similarly *in vivo*. Differences in flaviolin synthesis from varying coding sequences under pBAD expression are indistinguishable *in vivo* (**Figure 4C**), and generally lower compared to production in the cell-free system (**Figure 3C**). Efficient catabolism of L-arabinose by *E. coli* cells is a drawback of arabinose-inducible promoters, which may be a potential cause for better flaviolin synthesis in pBAD-regulated CFE. ^28^ Notably, the current CFE system cannot be used to prototype promoter choice for the *in vivo* expression of RppA, given any coding sequence (**Figure 3 and Figure 4**). The pT7 expression measurements were collected when the inducible IPTG was added, which was after 3.5 hours of growth (**Figure 4A**). While the measurement of the pJ23101 constructs were collected right after inoculation into the 96-well plate. It takes about 3-4 hours for the cells to grow before they start to produce pigment. This explains the delay occurring in the pJ23101 promoter (**Figure 4D**). This is also true for sfGFP expression. While sfGFP expressed highest using pT7 in both systems, the pBAD-sfGFP expressed better in CFE while the constitutive promoter and pTet promoter are expressed better *in vivo* (**Figures S7D&4E**). From this set of experiments, this suggests that there is strong correlation, but not complete alignment between expressing in cell free vs. *in vivo*.

**Figure 4.**
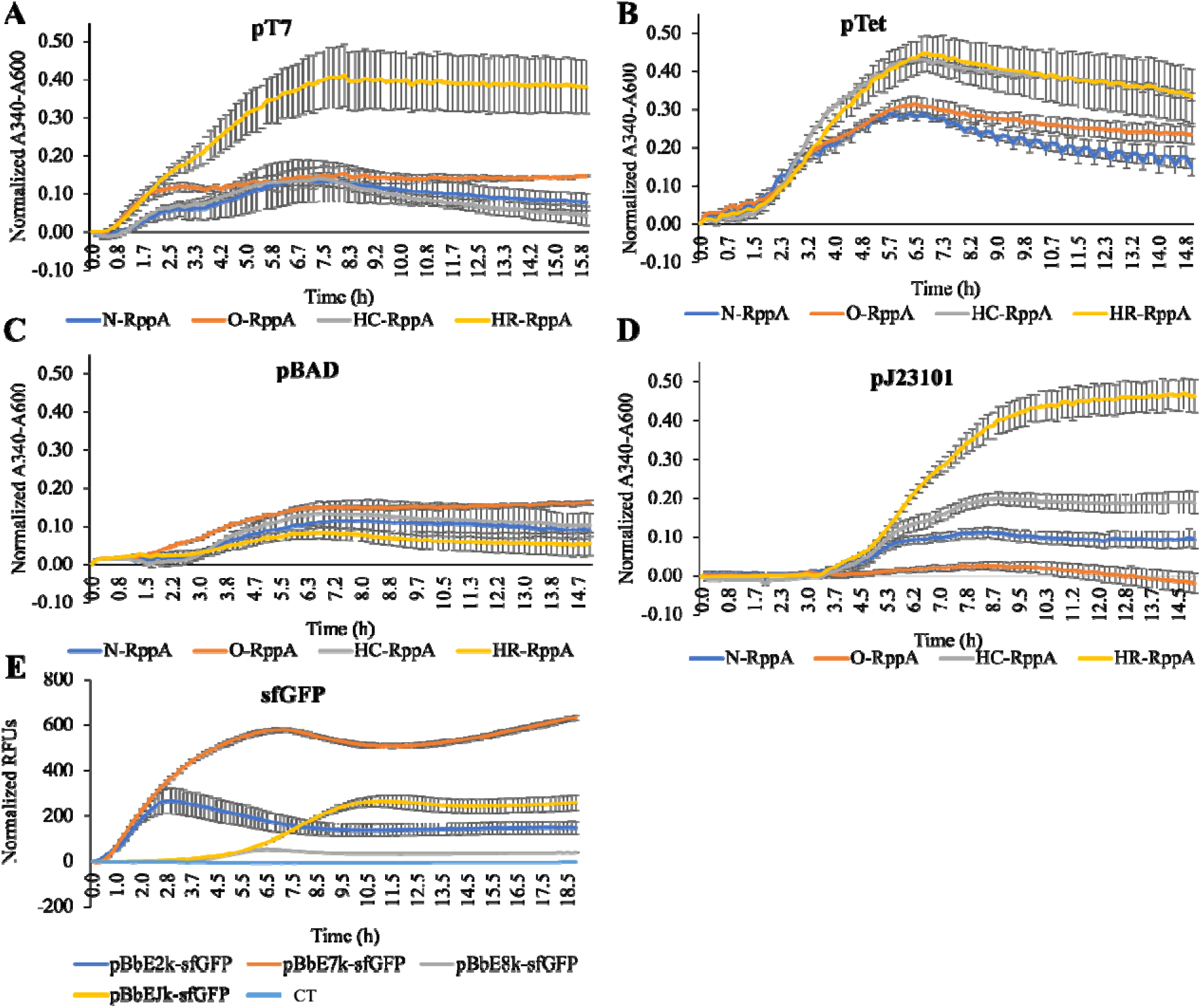
Flaviolin formation in *E. coli* BL21 (Star)DE3 cells carrying different promoter and coding sequence combinations. Flaviolin production in cells carrying (A) pT7 promoter constructs post-induction with 500 µM IPTG (added after 3.5 hours of growth), (B) pTet promoter constructs post-induction with100 nM anhydrotetracycline (added after 3 hours of growth), (C) pBAD constructs post-induction with 5000 µM L-arabinose (added after 1 hour and 10 min), and (D) pJ23101 constructs. (E) Expression of BL21 Star(DE3) cells harboring sfGFP. Reactions were run in triplicate and read every 10 minutes for 20 hours. Error bars represent the standard error of the mean (*n* = 3).

## Conclusion

Natural product synthesis in non-native contexts requires the successful translation and folding of biosynthetic genes. Refactoring choices to improve the heterologous expression and activity of these enzymes include the use of inducible or constitutive promoters and codon optimization/harmonization strategies. These elements are commonly explored for *in vivo* protein synthesis purposes, but long Design-Build-Test-Learn (DBTL) cycles associated with cellular engineering can be a limiter when evaluating gene refactoring strategies. Cell-free systems enable the accelerated testing of such tactics and thus enable the rapid optimization of refactoring choices for *in vitro* or *in vivo* expression. However, besides pT7 promoter systems and a selection of constitutive promoters, ^72^ other promoter systems have not been used extensively in CFE. Tools for codon harmonization, as opposed to codon optimization, have also not been considered for CFE. Thus, we aimed to explore whether inducible promoter systems and codon harmonization can benefit *in vitro* protein synthesis or be prototyped in CFE reactions for *in vivo* implementation.

In the cell-free context, we show that inducible pTet– and pBAD-regulated expression, while slower than pT7, allow higher yields of flaviolin. The same is true for constitutive expression. As codon harmonization algorithms are likely to more accurately measure the “tempo” of translation elongation. ^42,73^ We demonstrate that even in a gene with mostly efficient codons (74.5% rank 1, 21.7% rank 2; avg rank 1.33 based on ROC-SEMPPR), codon harmonization improves natural product synthesis more than uniform codon optimization across the cell-free and cellular environments considered here (7/8 cases). This is consistent with Keasling and coworkers recent report that in some, but not all heterologous hosts, codon harmonization can be superior to other codon optimization methods to express a type I polyketide synthase gene from actinomycete origin,^7^ providing further support to the notion that codon harmonization should be explored more generally to promote improved protein production from biosynthetic genes from Actinobacteria. Interestingly, we found that different harmonization methods do not work equally well for *rppA*. Consistent with the original protein coding sequence having few “slow” codons, which probably affect co-translational folding the most,^33^ in 6 out of the 8 cases evolutionarily based harmonization, i.e, fitting the evolutionarily based ROC-SEMPPR to the donor and host genomes to determine and replace based on individual codon ranks, performed substantially better than the window-based CHARMING approach (in one of the two other cases, the differences were almost indistinguishable). We also found that the choice of promoters influences the outcome of refactoring coding sequences. These interactions also vary between *in vitro* and *in vivo* reactions, particularly for non-pT7 inducible promoters, for which the relative activities of promoters and the synergies between promoter usage and coding sequences poorly correlate between cell-free and cell-based systems. In conclusion, refactoring promoters and/or coding sequences via CFE can be a valuable strategy to rapidly screen for catalytically functional production of enzymes from BCGs. This can in turn accelerate DBTL cycles to generate valuable metabolites.

## Materials and methods

### Strains and plasmids

*E. coli* BL21 Star(DE3) was purchased from New England Biosciences (Ipswich, Massachusetts, USA). *Streptomyces griseus* was purchased from Carolina (cat# 155705). The *rppA* gene from *S. griseus* was codon optimized using Integrated DNA Technology’s (IDT) codon optimization tool (https://HR.idtdna.com). *rppA* codon optimized and *rppA* codon harmonized with a C-terminus strep tag, TGGAGCCATCCGCAGTTCGAAAAA, were ordered from IDT (**Table S1**). BioBrick plasmids were obtained from Addgene (https://HR.addgene.org): pBbE2k (Plasmid #35324), pBbE7k (Plasmid #35315), pBbE8k (Plasmid #35270), and pJL1-sfGFP (Plasmid #102634) (**Table S2**). All of the constructs were cloned via Gibson assembly (New England Biolabs, part #E2611S). Primers used in this study were designed with the J5 algorithm and are listed in **Table S3**.^74^

### *In vivo* Flaviolin measurement

*E. coli* BL21 Star(DE3) was used as a host strain for *in vivo* expression of RppA. Cultures were grown in 2xYPTG media (10 g/L yeast extract, 7 g/L potassium phosphate dibasic, 3 g/L potassium phosphate monobasic, 5 g/L NaCl, 16 g/L tryptone, and 18 g/L glucose) supplemented with kanamycin at 50 µg/mL. Overnight seed cultures (2 mL) were grown from a fresh single colony at 37°C, shaking at 210 rpm. In a 96-well plate (Greiner), all constructs started growing with the initial OD_600_= 0.005. 5000 µM L-arabinose, 100 nM anhydrotetracycline, and 500 µM IPTG were induced after 110 min, 180 min, and 210 min, respectively. The plate was covered with an adhesive plate seal (Thermo Scientific) and loaded measured on a VARIOSKAN LUX (Thermo Scientific) plate reader. Readings at A_340_/A_600_ were taken every 10 min for 20 hr.

### *In vivo* Flaviolin Production and Purification

A 25 mL seed culture of BL21 Star(DE3) harboring the optimized *rppA* driven by pT7 plasmid was grown overnight (37 °C, 2010 rev/min) in LB medium supplemented with 50 µg/mL kanamycin. After ∼20 hours, 10 mL of seed culture was used to inoculate 1 L media in a Fernbach flask (VWR 29171-854). Cells were incubated at 37 °C shaking at 210 rev/min. At an OD_600_ of ∼0.8, the culture was induced with 0.5 mM IPTG and grown at 16 °C for 20 hr. The culture was then centrifuged at 5000 x g for 30 mins. The pink supernatant was adjusted to pH=2 with 3M HCl and incubated at 4°C overnight to precipitate flaviolin. Pigments were recovered by centrifugation at 5000 x g for 30 mins, and the precipitate was washed with DI water. The pellet was then dried at 50°C in an oven overnight. The dried pellets were washed with 6M HCl at 100°C to remove proteins and carbohydrates, then centrifuged at 5000 x g for 10 mins. The precipitate was washed with ethanol and chloroform and then dried at 50°C overnight. Purity was verified via Direct Analysis in Real Time Mass Spectrometry – DART-MS (*m/z* = 206), and ^1^H NMR: (400 mHz, acetone D_6_) δ 12.53 ppm (s, OH), 2.29 ppm (m, CH), 6.11 ppm (s, CH), 7.08 ppm (d, CH) and 6.61 ppm (d, CH) which is in agreement with the previous literature report.^75,76^

### Cell-free extract preparation

The same extract preparation procedure was used for all strains. A seed culture was prepared with 30 mL 2xYPTG media (10 g/L yeast extract, 7 g/L potassium phosphate dibasic, 3 g/L potassium phosphate monobasic, 5 g/L NaCl, 16 g/L tryptone, and 18 g/L glucose) inoculated with a fresh colony and incubated overnight at 37°C, 220 rpm. 1 L of 2xYPTG media in a 2.5 L Tunair flask was then inoculated with the overnight culture and grown at 37°C, 220 rpm. Cell growth was monitored by NanoDrop (Thermo Scientific). Cells were harvested at OD_600_ ∼ 2.8-3.2 by centrifugation (5000 x g, 15 min, 10°C), then washed three times using S30 buffer (10 mM Tris-acetate, 14 mM magnesium acetate, 60 mM potassium acetate, and 10 mM DTT). All wash steps were performed at 4°C. Cell pellets were then weighed, flash frozen, and then stored at –80°C. For extract preparation, the cell pellets were then thawed on ice and resuspended in 0.8mL of S30 buffer per g of cell pellet of the pellet by vortexing with short bursts (vortex 15 s, rest 30 s, repeat). 1.4 mL aliquots were sonicated on ice in 2 mL microcentrifuge tubes using an OMNI Sonic Rupto 400 (45 s on, 59 s off for three cycles, 50% amplitude set). 4.5 µL of 1 M DTT was added into each tube immediately after sonication. All samples were centrifuged at 12000 x g for 10 minutes at 4°C. The supernatant was collected without disturbing the pellet and centrifuged again to remove the remaining debris. The resulting supernatants were aliquoted into fresh centrifuge tubes, flash-frozen, and stored at –80°C.

### CFE reaction preparation

The cell-free reaction comprised 1.2 mM ATP, 0.85 mM GTP, 0.85 mM UTP and 0.85 mM CTP; 34.5 μg/mL folinic acid; 0.4 mM nicotinamide adenine dinucleotide (NAD), 0.27 mM coenzyme A (CoA), 4 mM oxalic acid, 1 mM, 1.5 mM spermidine, 57.33 mM HEPES buffer, 10 mM magnesium glutamate, 10 mM ammonium glutamate, 130 mM potassium glutamate, 2 mM each of the 20 amino acids, 33 mM phosphoenolpyruvate (PEP), 27 ng/µL DNA template and incubated at 30°C.^65^ Reactions were set up in 10 µL volumes unless otherwise stated. The type of inducer used and changes to any of these conditions are described in the text. All reactions were incubated in a 96 PCR well plate (VWR #47744-116). Surrounding wells were filled with 1x phosphate-buffered saline (PBS) to control the humidity and prevent evaporation. Plates were covered with an adhesive plate seal (Thermo Scientific), before putting it in the plate reader. Flaviolin synthesis was monitored by reading reaction absorbance at 340 nm at varying timeframes and intervals, as described in the text.

### Flaviolin quantitation in lysates with absorbance measurements

To generate standard curves from pigment absorbance measurements, increasing concentrations of the purified pigment dissolved in DMSO were spiked into BL21 Star(DE3) lysate mock reactions (i.e., reactions without DNA). Absorbance measurements were made in a 96 PCR well plate, without a lid, loaded into a VARIOSKAN LUX (Thermo Scientific) plate reader. The read protocol was set to shake the plate at high speed for 2 s then measure absorbance in selected wells at 340 nm. The resulting values were then normalized to the 0 µM pigment condition.

To measure the absorbance of flaviolin produced by cell-free expressed rppA, base reaction mixes with BL21 Star(DE3) lysate and RppA-expressing plasmid DNA were performed. Modifications to the reactions for validating pigment production are described in the text. All reactions were laid out on a 96-PCR well plate and measured every 10s for 20 HR at 30°C. A_340_ measurements were taken and normalized as described above.

### Quantification of active sfGFP

Fluorescence measurements of reactions expressing sfGFP were taken using top optics on a VARIOSKAN LUX (Thermo Scientific). Excitation and emission filters were set to 485 nm and 538 nm, respectively.

## Supplementary data

### Data availability

All the data described in this article are available within the article and in its online supplementary files.

## Funding

This work was supported by the University of Tennessee-Knoxville, the University of Tennessee-Oak Ridge Innovation Institute Science Alliance, the National Institutes of Health (R15GM145182), and the University of Sydney to C.B.B. C.B.B. is a member of the University of Sydney Drug Discovery Initiative, the University of Sydney Infectious Diseases Institute, and the University of Sydney Nanoscience Institute. This research was sponsored by the Genomic Science Program, US. Department of Energy, Office of Science, Biological, and Environmental Research as part of the Plant Microbe Interfaces Scientific Focus Area (http://pmi.ornl.gov). Oak Ridge National Laboratory is managed by UT-Battelle, LLC, for the Department of Energy under contract DE-AC05099OR2725. This manuscript has been authored by UT-Battelle, LLC under Contract DA-AC05-00OR2275 with the U.S. Department of Energy. The United States Government retains a nonexclusive, paid-up, irrevocable worldwide license to publish or reproduce the published form of this manuscript or allow others to do so, for United States Government purposes. The Department of Energy will provide public access to these results of federally sponsored research in accordance with the DOE Access plan (ttp://energy.gov/downloads/doe-public-access-plan). J.L.N.D. and T.T.S. were supported by University of Tennessee-Oak Ridge Innovation Institute Science Alliance Graduate-Advancement, Training, and Education (GATE) fellowships. G.A., J.W.B., D.S.G., and E.G., were supported by the Advanced Undergraduate Research Activity (AURA) fellowships from the University of Tennessee-Knoxville Office of Undergraduate Research and Fellowships. M.S. was supported by a Summer Research Training (SmART) summer internship from the UT-Oak Ridge Innovation Institute (UT-ORII).

## Conflict of interest statement

All the authors declare that they have no competing interests.

## Notes

### Competing Interest Statement

The authors have declared no competing interest.

