## Supplementary Information for "Profiling Expression Strategies for a Type III Polyketide Synthase in a Lysate-Based, Cell-free System"

**Table S1.** Genes sequences were used in this study; highlighted nucleotides are for the strep tag.

| Name | Sequence |
|---|---|
| Native *rppA* | atggcgaccctgtgccgaccggccatcgctgtgcccgagcacgtcatcacgatgcagcagaccctggacctg<br>gcccgggagacccatgccgggcacccgcagcgcgacctcgtcctgaggctcatccagaacaccggcgtcca<br>gacccggcacctcgtgcagcccatcgagaagaccctggcgcaccccggattcgaggtgcgcaaccaggtgta<br>cgaggccgaggccaagacccgggtccccgaagtcgtccggcgggcgctcgccaacgccgagaccgagcc<br>gtccgagatcgacctgatcgtctacgtctcctgcacgggtttcatgatgccctcgctgaccgcgtggatcatcaac<br>agcatgggcttccggcccgagacccgccaactgcccatcgcccagctcggctgtgcggcgggcggcgcggc<br>gatcaaccgcgcgcacgacttctgcgtggcctaccccgactccaacgtcctcatcgtgtcctgcgagttctgctc<br>gctgtgctaccagcccaccgacatcggggtcggttccctgctctccaacggactcttcggcgacgcgctctccg<br>cggccgtcgtacggggacagggcggcaccggcatgcgcctggagcgcaacggctcccacctggtgcccga<br>caccgaggactggatctcctacgcggtccgcgacaccgggttccacttccagctggacaagcgggtcccggg<br>caccatggagatgctcgccccggtgctcctggacctggtcgacctgcacggctggtccgtcccgaacatggac<br>ttcttcatcgtccacgcgggcggaccgcgcatcctggacgacctctgccacttcctcgacctgccgcccgagat<br>gttccgctacagccgggccaccctcaccgaacgcggcaacatcgcgagctccgtcgtcttcgacgcgctggc<br>gcgcctcttcgacgacggcggcgccgccgagtccgcgcaggggctcatcgccggcttcggtcccggcatcac<br>cgccgaggtggccgtggggagttgggccaaggaaggcctcggggcggacgtcggacgcgacctcgacga<br>gctggagctgaccgccggcgttgcgctgtccggctggagccacccgcagttcgaaaaataa |

| | |
|---|---|
| *E. coli* codon optimized *rppA* | atggctactctgtgtcgccctgcgattgccgttccagagcacgttatcacgatgcaacaaactttggatcttgcccg<br>tgagacccacgcaggccatcctcaacgtgacctggtgttacgtttgatccagaatactggagttcaaacccgcca<br>tttagttcagccaatcgaaaagacattggcacatcctgggtttgaagttcgcaaccaagtctatgaagctgaagca<br>aaaacgcgcgtacctgaagtggttcgccgtgctctggcgaatgcggaaacggagcccagtgaaatcgatttaat<br>tgtgtatgtttcttgcaccggttttatgatgccgtctcttacggcatggattatcaatagtatgggctttcgtcctgaga<br>cacgccaactgccaatcgcacaattaggatgtgcagcgggaggtgcggcgattaaccgcgcccatgatttttgc<br>gtagcataccccgactccaatgttttaattgtgagctgtgagttttgctcgctgtgttatcaaccaacggatattggc<br>gtgggttccttgttgtccaacggattgtttggggacgcccttagtgcggctgttgtgcgtggtcagggggtaccg<br>gcatgcgccttgaacgcaatgggtcccacttggttcccgacacggaggattggatctcatacgcagtgcgtgata<br>caggcttccatttccaattagacaaacgcgtacctgggacgatggaaatgttagcgccggtactgttggacctgg<br>tggatctgcacggttggagcgtgccaaatatggactttttatcgtgcacgccggtggacctcgcatccttgatga<br>cttatgtcacttttggatctgcctccagagatgttccgctactctcgcgccactcttacggaacgcggcaatattgc<br>aagctcagtcgtatttgatgcattggcgcgcttatttgacgacggaggcgcagccgagtccgcccaagggctga<br>tcgcggggtttggtccggggatcacggcggaagtagcagtcggtagctgggccaaggaggggttaggagcg<br>gacgttgggcgtgatctggatgagttggaactgaccgccggagttgctctgagtggatggagccacccgcagtt<br>cgaaaaataa |
| *rppA* codon harmonized using the CHARMING algorithm | atggccaccctgtgcagaccggcgattgctgttccagaacatgtgattacgatgcagcagaccctggatctggc<br>ccgtgaaacccacgcgggtcatccgcagcgcgatttagtgctgcgtctgattcagaacaccggcgtgcagacc<br>cgtcatctggttcagccaattgaaaaaaccctggcccatccaggatttgaagtgcgcaaccaggtttatgaagcg<br>gaagcgaaaacccgcgtgccagaggtggtgcgccgcgcgctggccaacgcggaaaccgaaccgagcgaa<br>attgatctgattgtgtatgtgagctgcacggggtttatgatgccaagcctgaccgcctggattattaactccatggg<br>ctttcgtccggaaacccgccaactgccgattgcccagctgggctgtgccgccggtggtgccgccattaaccgc<br>gcccatgatttttgcgttgcgtatccggatagcaacgtgctgattgttagttgcgaattttgcagtctgtgctatcagc<br>caaccgatattggtgtggggagcctgttaagcaacggtttatttggcgatgcgctgagcgccgcggtggtacgt<br>ggacagggcggcaccggcatgcgcctggaacgcaacggcagccatctggtgccagataccgaagattggatt<br>agctatgccgttcgcgataccggttttcactttcagctggataaacgtgtgccgggcaccatggaaatgctggcc<br>ccggtgctgctggacctggtggatctgcatggctggagcgtgccgaacatggattttttattgtgcatgccggcg<br>gaccgcgcattctggatgatttatgccattttctggatctgccgccagaaatgtttcgctattcccgtgcgaccctga<br>ccgagcgcggcaacattgcctccagcgtggtgtttgatgcactggcacgtctgttcgatgatggcggcgcggc<br>ggaaagcgcccagggcttaattgcgggctttggtccaggcattaccgcggaagttgcggttggttcgtgggcga<br>aagaaggcttaggtgccgatgtgggacgcgatctggatgaactggaactgacggcgggcgtagccctgagcg<br>gctggagccacccgcagttcgaaaaatag |
| *rppA* codon harmonized using the ROC-SEMPPR algorithm | atggcaactctctgccggcctgctatcgccgtgccggaacacgttatcaccatgcagcagactctcgacctcgct<br>cgcgaaactcatgctgggcaccctcagcgtgacctggttctccgactgatccagaacactggtgttcagactcgc<br>cacctggtgcagccgatcgaaaaaactctcgcacacccgggcttcgaagtgcgtaaccaggtgtacgaagctg<br>aagctaaaactcgcgttccggaggttgttcgccgcgcactggctaacgctgaaactgaaccttctgaaatcgacc<br>tcatcgtttacgtttcttgcaccggattcatgatgccgtccctcactgcatggatcatcaacagcatgggtttccgcc<br>cggaaactcgtcaactcccgatcgctcagctgggttgtgcagcaggtggtgcagcaatcaaccgtgcacacga<br>cttctgcgtggcttacccggactctaacgttctgatcgtgtcttgcgaattctgctccctctgctaccagccgactga<br>catcggggttggatctctcctgtctaacggcctgttcggtgacgcactgtctgcagctgttgtacgcggccagggt<br>ggtactggtatgcgtctcgaacgtaacggttctcacctcgtgccggacactgaagactggatctcttacgcagttc<br>gtgacactgggttccacttccagctcgacaaacgcgttcctggtactatggaaatgctggctcctgtgctgctcga<br>cctcgttgacctccacggttggtctgttcctaacatggacttcttcatcgttcacgcaggtggccctcgtatcctcga<br>cgacctgtgccacttcctggacctccctccggaaatgttccgttacagccgcgctactctgactgagcgtggtaa<br>catcgcaagctctgttgttttcgacgcactcgcacgtctgttcgacgacggtggtgctgctgaatctgcacagggg<br>ctgatcgctggtttcggaccgggtatcactgctgaagtggctgtggggagttgggctaaagagggtctgggggc |

| | |
|---|---|
| | agacgttggccgtgacctggacgaactcgaactcactgctggtgtcgcactctctggttggagccacccgcagtt<br>cgaaaaataa |
| *sf*GFP | atgagcaaaggtgaagaactgtttaccggcgttgtgccgattctggtggaactggatggcgatgtgaacggtca<br>caaattcagcgtgcgtggtgaaggtgaaggcgatgccacgattggcaaactgacgctgaaatttatctgcacca<br>ccggcaaactgccggtgccgtggccgacgctggtgaccaccctgacctatggcgttcagtgttttagtcgctatc<br>cggatcacatgaaacgtcacgatttctttaaatctgcaatgccggaaggctatgtgcaggaacgtacgattagcttt<br>aaagatgatggcaaatataaaacgcgcgccgttgtgaaatttgaaggcgataccctggtgaaccgcattgaact<br>gaaaggcacggattttaaagaagatggcaatatcctgggccataaactggaatacaactttaatagccataatgtt<br>tatattacggcggataaacagaaaaatggcatcaaagcgaattttaccgttcgccataacgttgaagatggcagt<br>gtgcagctggcagatcattatcagcagaataccccgattggtgatggtccggtgctgctgccggataatcattatc<br>tgagcacgcagaccgttctgtctaaagatccgaacgaaaaaggcacgcgggaccacatggttctgcacgaata<br>tgtgaatgcggcaggtattacgtggagccatccgcagttcgaaaaataa |

**Table S2.** Plasmid sequences were used in this study; highlighted nucleotides are for the promoters.

| Name | Sequence |
|---|---|
| pBbE2k-containing pTet promoter | ggatccaaactcgagtaaggatctccaggcatcaaataaaacgaaaggctcagtcgaaagactgggcctttcgt<br>tttatctgttgtttgtcggtgaacgctctctactagagtcacactggctcaccttcgggtgggcctttctgcgtttatac<br>ctagggcgttcggctgcggcgagcggtatcagctcactcaaaggcggtaatacggttatccacagaatcaggg<br>gataacgcaggaaagaacatgtgagcaaaaggccagcaaaaggccaggaaccgtaaaaaggccgcgttgct<br>ggcgtttttccataggctccgcccccctgacgagcatcacaaaaatcgacgctcaagtcagaggtggcgaaacc<br>cgacaggactataaagataccaggcgtttccccctggaagctccctcgtgcgctctcctgttccgaccctgccgc<br>ttaccggatacctgtccgcctttctcccttcgggaagcgtggcgctttctcatagctcacgctgtaggtatctcagtt<br>cggtgtaggtcgttcgctccaagctgggctgtgtgcacgaaccccccgttcagcccgaccgctgcgccttatcc<br>ggtaactatcgtcttgagtccaacccggtaagacacgacttatcgccactggcagcagccactggtaacaggatt<br>agcagagcgaggtatgtaggcggtgctacagagttcttgaagtggtggcctaactacggctacactagaagga<br>cagtatttggtatctgcgctctgctgaagccagttaccttcggaaaaagagttggtagctcttgatccggcaaaca<br>aaccaccgctggtagcggtggtttttttgtttgcaagcagcagattacgcgcagaaaaaaaggatctcaagaaga<br>tcctttgatcttttctacggggtctgacgctcagtggaacgaaaactcacgttaagggattttggtcatgactagtgc<br>ttggattctcaccaataaaaaacgcccggcggcaaccgagcgttctgaacaaatccagatggagttctgaggtc<br>attactggatctatcaacaggagtccaagcgagctctcgaaccccagagtcccgctcagaagaactcgtcaaga<br>aggcgatagaaggcgatgcgctgcgaatcgggagcggcgataccgtaaagcacgaggaagcggtcagccc<br>attcgccgccaagctcttcagcaatatcacgggtagccaacgctatgtcctgatagcggtccgccacacccagc<br>cggccacagtcgatgaatccagaaaagcggccattttccaccatgatattcggcaagcaggcatcgccatgggt<br>cacgacgagatcctcgccgtcgggcatgcgcgccttgagcctggcgaacagttcggctggcgcgagcccctg<br>atgctcttcgtccagatcatcctgatcgacaagaccggcttccatccgagtacgtgctcgctcgatgcgatgtttcg<br>cttggtggtcgaatgggcaggtagccggatcaagcgtatgcagccgccgcattgcatcagccatgatggatact<br>ttctcggcaggagcaaggtgagatgacaggagatcctgccccggcacttcgcccaatagcagccagtcccttc<br>ccgcttcagtgacaacgtcgagcacagctgcgcaaggaacgcccgtcgtggccagccacgatagccgcgctg<br>cctcgtcctgcagttcattcagggcaccggacaggtcggtcttgacaaaaagaaccgggcgcccctgcgctga<br>cagccggaacacggcggcatcagagcagccgattgtctgttgtgcccagtcatagccgaatagcctctccacc<br>caagcggccggagaacctgcgtgcaatccatcttgttcaatcatgcgaaacgatcctcatcctgtctcttgatcag<br>atcatgatcccctgcgccatcagatccttggcggcaagaaagccatccagtttactttgcagggcttcccaacctt<br>accagagggcgccccagctggcaattccgacgtcttaagacccactttcacatttaagttgtttttctaatccgcat |

| | |
|---|---|
| | atgatcaattcaaggccgaataagaaggctggctctgcaccttggtgatcaaataattcgatagcttgtcgtaataa<br>tggcggcatactatcagtagtaggtgtttccctttcttctttagcgacttgatgctcttgatcttccaatacgcaaccta<br>aagtaaaatgccccacagcgctgagtgcatataatgcattctctagtgaaaaaccttgttggcataaaaaggctaa<br>ttgattttcgagagtttcatactgtttttctgtaggccgtgtacctaaatgtacttttgctccatcgcgatgacttagtaa<br>agcacatctaaaacttttagcgttattacgtaaaaaatcttgccagctttccccttctaaagggcaaaagtgagtatg<br>gtgcctatctaacatctcaatggctaaggcgtcgagcaaagcccgcttattttttacatgccaatacaatgtaggct<br>gctctacacctagcttctgggcgagtttacgggttgttaaaccttcgattccgacctcattaagcagctctaatgcg<br>ctgttaatcactttacttttatctaatctagacatcattaattcctaatttttgttgacactctatcgttgatagagttatttta<br>ccactccctatcagtgatagagaaaagaattcaaaagatcttttaagaaggagatatacat |
| pBbE7k-<br>containing<br>pT7<br>promoter | ggatccaaactcgagtaaggatctccaggcatcaaataaaacgaaaggctcagtcgaaagactgggcctttcgt<br>tttatctgttgtttgtcggtgaacgctctctactagagtcacactggctcaccttcgggtgggcctttctgcgtttatac<br>ctagggcgttcggctgcggcgagcggtatcagctcactcaaaggcggtaatacggttatccacagaatcaggg<br>gataacgcaggaaagaacatgtgagcaaaaggccagcaaaaggccaggaaccgtaaaaaggccgcgttgct<br>ggcgtttttccataggctccgcccccctgacgagcatcacaaaaatcgacgctcaagtcagaggtggcgaaacc<br>cgacaggactataaagataccaggcgtttccccctggaagctccctcgtgcgctctcctgttccgaccctgccgc<br>ttaccggatacctgtccgcctttctcccttcgggaagcgtggcgctttctcatagctcacgctgtaggtatctcagtt<br>cggtgtaggtcgttcgctccaagctgggctgtgtgcacgaaccccccgttcagcccgaccgctgcgccttatcc<br>ggtaactatcgtcttgagtccaacccggtaagacacgacttatcgccactggcagcagccactggtaacaggatt<br>agcagagcgaggtatgtaggcggtgctacagagttcttgaagtggtggcctaactacggctacactagaagga<br>cagtatttggtatctgcgctctgctgaagccagttaccttcggaaaaagagttggtagctcttgatccggcaaaca<br>aaccaccgctggtagcggtggtttttttgtttgcaagcagcagattacgcgcagaaaaaaaggatctcaagaaga<br>tcctttgatcttttctacggggtctgacgctcagtggaacgaaaactcacgttaagggattttggtcatgactagtgc<br>ttggattctcaccaataaaaaacgcccggcggcaaccgagcgttctgaacaaatccagatggagttctgaggtc<br>attactggatctatcaacaggagtccaagcgagctctcgaaccccagagtcccgctcagaagaactcgtcaaga<br>aggcgatagaaggcgatgcgctgcgaatcgggagcggcgataccgtaaagcacgaggaagcggtcagccc<br>attcgccgccaagctcttcagcaatatcacgggtagccaacgctatgtcctgatagcggtccgccacacccagc<br>cggccacagtcgatgaatccagaaaagcggccattttccaccatgatattcggcaagcaggcatcgccatgggt<br>cacgacgagatcctcgccgtcgggcatgcgcgccttgagcctggcgaacagttcggctggcgcgagcccctg<br>atgctcttcgtccagatcatcctgatcgacaagaccggcttccatccgagtacgtgctcgctcgatgcgatgtttcg<br>cttggtggtcgaatgggcaggtagccggatcaagcgtatgcagccgccgcattgcatcagccatgatggatact<br>ttctcggcaggagcaaggtgagatgacaggagatcctgccccggcacttcgcccaatagcagccagtcccttc<br>ccgcttcagtgacaacgtcgagcacagctgcgcaaggaacgcccgtcgtggccagccacgatagccgcgctg<br>cctcgtcctgcagttcattcagggcaccggacaggtcggtcttgacaaaaagaaccgggcgcccctgcgctga<br>cagccggaacacggcggcatcagagcagccgattgtctgttgtgcccagtcatagccgaatagcctctccacc<br>caagcggccggagaacctgcgtgcaatccatcttgttcaatcatgcgaaacgatcctcatcctgtctcttgatcag<br>atcatgatcccctgcgccatcagatccttggcggcaagaaagccatccagtttactttgcagggcttcccaacctt<br>accagagggcgccccagctggcaattccgacgtcctcactgcccgctttccagtcgggaaacctgtcgtgcca<br>gctgcattaatgaatcggccaacgcgcggggagaggcggtttgcgtattgggcgccagggtggtttttcttttca<br>ccagtgagacgggcaacagctgattgcccttcaccgcctggccctgagagagttgcagcaagcggtccacgct<br>ggtttgccccagcaggcgaaaatcctgtttgatggtggttaacggcgggatataacatgagctgtcttcggtatcg<br>tcgtatcccactaccgagatgtccgcaccaacgcgcagcccggactcggtaatggcgcgcattgcgcccagcg<br>ccatctgatcgttggcaaccagcatcgcagtgggaacgatgccctcattcagcatttgcatggtttgttgaaaacc<br>ggacatggcactccagtcgccttcccgttccgctatcggctgaatttgattgcgagtgagatatttatgccagcca<br>gccagacgcagacgcgccgagacagaacttaatgggcccgctaacagcgcgatttgctggtgacccaatgcg<br>accagatgctccacgcccagtcgcgtaccgtcttcatgggagaaaataatactgttgatgggtgtctggtcagag<br>acatcaagaaataacgccggaacattagtgcaggcagcttccacagcaatggcatcctggtcatccagcggata |

| | |
|---|---|
| | gttaatgatcagcccactgacgcgttgcgcgagaagattgtgcaccgccgctttacaggcttcgacgccgcttcg<br>ttctaccatcgacaccaccacgctggcacccagttgatcggcgcgagatttaatcgccgcgacaatttgcgacg<br>gcgcgtgcagggccagactggaggtggcaacgccaatcagcaacgactgtttgcccgccagttgttgtgccac<br>gcggttgggaatgtaattcagctccgccatcgccgcttccactttttcccgcgttttcgcagaaacgtggctggcct<br>ggttcaccacgcgggaaacggtctgataagagacaccggcatactctgcgacatcgtataacgttactggtttca<br>cattcaccaccctgaattgactctcttccgggcgctatcatgccataccgcgaaaggttttgcgccattcgatggtg<br>tccgggatctcgacgctctcccttatgcgactcctgcattaggaagcagcccagtagtaggttgaggccgttgag<br>caccgccgccgcaaggaatggtgcatgcaaggagatggcgcccaacagtcccccggccacggggcctgcc<br>accatacccacgccgaaacaagcgctcatgagcccgaagtggcgagcccgatcttccccatcggtgatgtcgg<br>cgatataggcgccagcaaccgcacctgtggcgccggtgatgccggccacgatgcgtccggcgtagaggatc<br>gagatcgatctcgatcccgcgaaattaatacgactcactatagggaattgtgagcggataacaatttcagaattc<br>aaaagatcttttaagaaggagatatacat |
| pBbE8k-containing pBAD promoter | ggatccaaactcgagtaaggatctccaggcatcaaataaaacgaaaggctcagtcgaaagactgggcctttcgt<br>tttatctgttgtttgtcggtgaacgctctctactagagtcacactggctcaccttcgggtgggcctttctgcgtttatac<br>ctagggcgttcggctgcggcgagcggtatcagctcactcaaaggcggtaatacggttatccacagaatcaggg<br>gataacgcaggaaagaacatgtgagcaaaaggccagcaaaaggccaggaaccgtaaaaaggccgcgttgct<br>ggcgtttttccataggctccgcccccctgacgagcatcacaaaaatcgacgctcaagtcagaggtggcgaaacc<br>cgacaggactataaagataccaggcgtttccccctggaagctccctcgtgcgctctcctgttccgaccctgccgc<br>ttaccggatacctgtccgcctttctcccttcgggaagcgtggcgctttctcatagctcacgctgtaggtatctcagtt<br>cggtgtaggtcgttcgctccaagctgggctgtgtgcacgaaccccccgttcagcccgaccgctgcgccttatcc<br>ggtaactatcgtcttgagtccaacccggtaagacacgacttatcgccactggcagcagccactggtaacaggatt<br>agcagagcgaggtatgtaggcggtgctacagagttcttgaagtggtggcctaactacggctacactagaagga<br>cagtatttggtatctgcgctctgctgaagccagttaccttcggaaaaagagttggtagctcttgatccggcaaaca<br>aaccaccgctggtagcggtggtttttttgtttgcaagcagcagattacgcgcagaaaaaaaggatctcaagaaga<br>tcctttgatcttttctacggggtctgacgctcagtggaacgaaaactcacgttaagggattttggtcatgactagtgc<br>ttggattctcaccaataaaaaacgcccggcggcaaccgagcgttctgaacaaatccagatggagttctgaggtc<br>attactggatctatcaacaggagtccaagcgagctctcgaaccccagagtcccgctcagaagaactcgtcaaga<br>aggcgatagaaggcgatgcgctgcgaatcgggagcggcgataccgtaaagcacgaggaagcggtcagccc<br>attcgccgccaagctcttcagcaatatcacgggtagccaacgctatgtcctgatagcggtccgccacacccagc<br>cggccacagtcgatgaatccagaaaagcggccattttccaccatgatattcggcaagcaggcatcgccatgggt<br>cacgacgagatcctcgccgtcgggcatgcgcgccttgagcctggcgaacagttcggctggcgcgagcccctg<br>atgctcttcgtccagatcatcctgatcgacaagaccggcttccatccgagtacgtgctcgctcgatgcgatgtttcg<br>cttggtggtcgaatgggcaggtagccggatcaagcgtatgcagccgccgcattgcatcagccatgatggatact<br>ttctcggcaggagcaaggtgagatgacaggagatcctgccccggcacttcgcccaatagcagccagtcccttc<br>ccgcttcagtgacaacgtcgagcacagctgcgcaaggaacgcccgtcgtggccagccacgatagccgcgctg<br>cctcgtcctgcagttcattcagggcaccggacaggtcggtcttgacaaaaagaaccgggcgcccctgcgctga<br>cagccggaacacggcggcatcagagcagccgattgtctgttgtgcccagtcatagccgaatagcctctccacc<br>caagcggccggagaacctgcgtgcaatccatcttgttcaatcatgcgaaacgatcctcatcctgtctcttgatcag<br>atcatgatcccctgcgccatcagatccttggcggcaagaaagccatccagtttactttgcagggcttcccaacctt<br>accagagggcgccccagctggcaattccgacgtcttatgacaacttgacggctacatcattcactttttcttcacaa<br>ccggcacggaactcgctcgggctggccccggtgcattttttaaatacccgcgagaaatagagttgatcgtcaaa<br>accaacattgcgaccgacggtggcgataggcatccgggtggtgctcaaaagcagcttcgcctggctgatacgtt<br>ggtcctcgcgccagcttaagacgctaatccctaactgctggcggaaaagatgtgacagacgcgacggcgaca<br>agcaaacatgctgtgcgacgctggcgatatcaaaattgctgtctgccaggtgatcgctgatgtactgacaagcct<br>cgcgtacccgattatccatcggtggatggagcgactcgttaatcgcttccatgcgccgcagtaacaattgctcaa<br>gcagatttatcgccagcagctccgaatagcgcccttccccttgcccggcgttaatgatttgcccaaacaggtcgct |

| | |
|---|---|
| | gaaatgcggctggtgcgcttcatccgggcgaaagaaccccgtattggcaaatattgacggccagttaagccatt<br>catgccagtaggcgcgcggacgaaagtaaacccactggtgataccattcgcgagcctccggatgacgaccgt<br>agtgatgaatctctcctggcgggaacagcaaaatatcacccggtcggcaaacaaattctcgtccctgatttttcac<br>caccccctgaccgcgaatggtgagattgagaatataacctttcattcccagcggtcggtcgataaaaaaatcgag<br>ataaccgttggcctcaatcggcgttaaacccgccaccagatgggcattaaacgagtatcccggcagcagggga<br>tcattttgcgcttcagccatacttttcatactcccgccattcagagaagaaaccaattgtccatattgcatcagacatt<br>gccgtcactgcgtcttttactggctcttctcgctaaccaaaccggtaaccccgcttattaaaagcattctgtaacaaa<br>gcgggaccaaagccatgacaaaaacgcgtaacaaaagtgtctataatcacggcagaaaagtccacattgattat<br>ttgcacggcgtcacactttgctatgccatagcatttttatccataagattagcggattctacctgacgcttttttatcgca<br>actctctactgtttctccatacccgtttttttgggaattcaaaagatcttttaagaaggagatatacat |
| pBbEJ23101<br>k-containing<br>pJ23101<br>promoter | tggagccacccgcagttcgaaaaataaggatccaaactcgagtaaggatctccaggcatcaaataaaacgaaa<br>ggctcagtcgaaagactgggcctttcgttttatctgttgtttgtcggtgaacgctctctactagagtcacactggctc<br>accttcgggtgggcctttctgcgtttatacctagggcgttcggctgcggcgagcggtatcagctcactcaaaggc<br>ggtaatacggttatccacagaatcaggggataacgcaggaaagaacatgtgagcaaaaggccagcaaaaggc<br>caggaaccgtaaaaaggccgcgttgctggcgtttttccataggctccgcccccctgacgagcatcacaaaaatc<br>gacgctcaagtcagaggtggcgaaacccgacaggactataaagataccaggcgtttccccctggaagctccct<br>cgtgcgctctcctgttccgaccctgccgcttaccggatacctgtccgcctttctcccttcgggaagcgtggcgcttt<br>ctcatagctcacgctgtaggtatctcagttcggtgtaggtcgttcgctccaagctgggctgtgtgcacgaacccc<br>cgttcagcccgaccgctgcgccttatccggtaactatcgtcttgagtccaacccggtaagacacgacttatcgcc<br>actggcagcagccactggtaacaggattagcagagcgaggtatgtaggcggtgctacagagttcttgaagtggt<br>ggcctaactacggctacactagaaggacagtatttggtatctgcgctctgctgaagccagttaccttcggaaaaa<br>gagttggtagctcttgatccggcaaacaaaccaccgctggtagcggtggtttttttgtttgcaagcagcagattac<br>gcgcagaaaaaaaggatctcaagaagatcctttgatcttttctacggggtctgacgctcagtggaacgaaaactc<br>acgttaagggattttggtcatgactagtgcttggattctcaccaataaaaaacgcccggcggcaaccgagcgttct<br>gaacaaatccagatggagttctgaggtcattactggatctatcaacaggagtccaagcgagctctcgaacccca<br>gagtcccgctcagaagaactcgtcaagaaggcgatagaaggcgatgcgctgcgaatcgggagcggcgatac<br>cgtaaagcacgaggaagcggtcagcccattcgccgccaagctcttcagcaatatcacgggtagccaacgctat<br>gtcctgatagcggtccgccacacccagccggccacagtcgatgaatccagaaaagcggccattttccaccatg<br>atattcggcaagcaggcatcgccatgggtcacgacgagatcctcgccgtcgggcatgcgcgccttgagcctgg<br>cgaacagttcggctggcgcgagcccctgatgctcttcgtccagatcatcctgatcgacaagaccggcttccatcc<br>gagtacgtgctcgctcgatgcgatgtttcgcttggtggtcgaatgggcaggtagccggatcaagcgtatgcagc<br>cgccgcattgcatcagccatgatggatactttctcggcaggagcaaggtgagatgacaggagatcctgccccg<br>gcacttcgcccaatagcagccagtcccttcccgcttcagtgacaacgtcgagcacagctgcgcaaggaacgcc<br>cgtcgtggccagccacgatagccgcgctgcctcgtcctgcagttcattcagggcaccggacaggtcggtcttga<br>caaaaagaaccgggcgcccctgcgctgacagccggaacacggcggcatcagagcagccgattgtctgttgtg<br>cccagtcatagccgaatagcctctccacccaagcggccggagaacctgcgtgcaatccatcttgttcaatcatgc<br>gaaacgatcctcatcctgtctcttgatcagatcatgatcccctgcgccatcagatccttggcggcaagaaagccat<br>ccagtttactttgcagggcttcccaaccttaccagagggcgccccagctggcaattccgacgtctttacagctagc<br>tcagtcctaggtattatgctagcaagaattcaaaagatcttttaagaaggagatatacat |
| pET28b-<br>containing<br>pT7<br>promoter | caccaccactgagatccggctgctaacaaagcccgaaaggaagctgagttggctgctgccaccgctgagcaat<br>aactagcataaccccttggggcctctaaacgggtcttgaggggttttttgctgaaaggaggaactatatccggatt<br>ggcgaatgggacgcgccctgtagcggcgcattaagcgcggcgggtgtggtggttacgcgcagcgtgaccgc<br>tacacttgccagcgccctagcgcccgctcctttcgctttcttcccttcctttctcgccacgttcgccggctttccccgt<br>caagctctaaatcgggggctccctttagggttccgatttagtgctttacggcacctcgaccccaaaaaacttgatta<br>gggtgatggttcacgtagtgggccatcgccctgatagacggtttttcgccctttgacgttggagtccacgttcttta<br>atagtggactcttgttccaaactggaacaacactcaaccctatctcggtctattcttttgatttataagggattttgccg |

| | |
|---|---|
| | atttcggcctattggttaaaaaatgagctgatttaacaaaaatttaacgcgaattttaacaaaatattaacgtttacaat<br>ttcaggtggcacttttcggggaaatgtgcgcggaacccctatttgtttatttttctaaatacattcaaatatgtatccgc<br>tcatgaattaattcttagaaaaactcatcgagcatcaaatgaaactgcaatttattcatatcaggattatcaataccat<br>atttttgaaaaagccgtttctgtaatgaaggagaaaactcaccgaggcagttccataggatggcaagatcctggta<br>tcggtctgcgattccgactcgtccaacatcaatacaacctattaatttcccctcgtcaaaaataaggttatcaagtga<br>gaaatcaccatgagtgacgactgaatccggtgagaatggcaaaagtttatgcatttctttccagacttgttcaacag<br>gccagccattacgctcgtcatcaaaatcactcgcatcaaccaaaccgttattcattcgtgattgcgcctgagcgag<br>acgaaatacgcgatcgctgttaaaaggacaattacaaacaggaatcgaatgcaaccggcgcaggaacactgcc<br>agcgcatcaacaatattttcacctgaatcaggatattcttctaatacctggaatgctgttttcccggggatcgcagtg<br>gtgagtaaccatgcatcatcaggagtacggataaaatgcttgatggtcggaagaggcataaattccgtcagcca<br>gtttagtctgaccatctcatctgtaacatcattggcaacgctacctttgccatgtttcagaaacaactctggcgcatc<br>gggcttcccatacaatcgatagattgtcgcacctgattgcccgacattatcgcgagcccatttatacccatataaat<br>cagcatccatgttggaatttaatcgcggcctagagcaagacgtttcccgttgaatatggctcataacaccccttgta<br>ttactgtttatgtaagcagacagttttattgttcatgaccaaaatcccttaacgtgagttttcgttccactgagcgtcag<br>accccgtagaaaagatcaaaggatcttcttgagatcctttttttctgcgcgtaatctgctgcttgcaaacaaaaaaac<br>caccgctaccagcggtggtttgtttgccggatcaagagctaccaactctttttccgaaggtaactggcttcagcag<br>agcgcagataccaaatactgtccttctagtgtagccgtagttaggccaccacttcaagaactctgtagcaccgcct<br>acatacctcgctctgctaatcctgttaccagtggctgctgccagtggcgataagtcgtgtcttaccgggttggactc<br>aagacgatagttaccggataaggcgcagcggtcgggctgaacggggggttcgtgcacacagcccagcttgga<br>gcgaacgacctacaccgaactgagatacctacagcgtgagctatgagaaagcgccacgcttcccgaagggag<br>aaaggcggacaggtatccggtaagcggcagggtcggaacaggagagcgcacgagggagcttccagggggg<br>aaacgcctggtatctttatagtcctgtcgggtttcgccacctctgacttgagcgtcgatttttgtgatgctcgtcagg<br>ggggcggagcctatggaaaaacgccagcaacgcggcctttttacggttcctggccttttgctggccttttgctcac<br>atgttctttcctgcgttatcccctgattctgtggataaccgtattaccgcctttgagtgagctgataccgctcgccgca<br>gccgaacgaccgagcgcagcgagtcagtgagcgaggaagcggaagagcgcctgatgcggtattttctcctta<br>cgcatctgtgcggtatttcacaccgcatatatggtgcactctcagtacaatctgctctgatgccgcatagttaagcc<br>agtatacactccgctatcgctacgtgactgggtcatggctgcgccccgacacccgccaacacccgctgacgcg<br>ccctgacgggcttgtctgctcccggcatccgcttacagacaagctgtgaccgtctccgggagctgcatgtgtcag<br>aggttttcaccgtcatcaccgaaacgcgcgaggcagctgcggtaaagctcatcagcgtggtcgtgaagcgattc<br>acagatgtctgcctgttcatccgcgtccagctcgttgagtttctccagaagcgttaatgtctggcttctgataaagcg<br>ggccatgttaagggcggttttttcctgtttggtcactgatgcctccgtgtaagggggatttctgttcatgggggtaat<br>gataccgatgaaacgagagaggatgctcacgatacgggttactgatgatgaacatgcccggttactggaacgtt<br>gtgagggtaaacaactggcggtatggatgcggcgggaccagagaaaaatcactcagggtcaatgccagcgct<br>tcgttaatacagatgtaggtgttccacagggtagccagcagcatcctgcgatgcagatccggaacataatggtgc<br>agggcgctgacttccgcgtttccagactttacgaaacacggaaaccgaagaccattcatgttgttgctcaggtcg<br>cagacgttttgcagcagcagtcgcttcacgttcgctcgcgtatcggtgattcattctgctaaccagtaaggcaacc<br>ccgccagcctagccgggtcctcaacgacaggagcacgatcatgcgcacccgtggggccgccatgccggcga<br>taatggcctgcttctcgccgaaacgtttggtggcgggaccagtgacgaaggcttgagcgagggcgtgcaagat<br>tccgaataccgcaagcgacaggccgatcatcgtcgcgctccagcgaaagcggtcctcgccgaaaatgaccca<br>gagcgctgccggcacctgtcctacgagttgcatgataaagaagacagtcataagtgcggcgacgatagtcatg<br>ccccgcgcccaccggaaggagctgactgggttgaaggctctcaagggcatcggtcgagatcccggtgcctaa<br>tgagtgagctaacttacattaattgcgttgcgctcactgcccgctttccagtcgggaaacctgtcgtgccagctgc<br>attaatgaatcggccaacgcgcggggagaggcggtttgcgtattgggcgccagggtggtttttcttttcaccagt<br>gagacgggcaacagctgattgcccttcaccgcctggccctgagagagttgcagcaagcggtccacgctggttt<br>gccccagcaggcgaaaatcctgtttgatggtggttaacggcgggatataacatgagctgtcttcggtatcgtcgta<br>tcccactaccgagatatccgcaccaacgcgcagcccggactcggtaatggcgcgcattgcgcccagcgccat |

| | |
|---|---|
| | ctgatcgttggcaaccagcatcgcagtgggaacgatgccctcattcagcatttgcatggtttgttgaaaaccggac<br>atggcactccagtcgccttcccgttccgctatcggctgaatttgattgcgagtgagatatttatgccagccagcca<br>gacgcagacgcgccgagacagaacttaatgggcccgctaacagcgcgatttgctggtgacccaatgcgacca<br>gatgctccacgcccagtcgcgtaccgtcttcatgggagaaaataatactgttgatgggtgtctggtcagagacat<br>caagaaataacgccggaacattagtgcaggcagcttccacagcaatggcatcctggtcatccagcggatagtta<br>atgatcagcccactgacgcgttgcgcgagaagattgtgcaccgccgctttacaggcttcgacgccgcttcgttct<br>accatcgacaccaccacgctggcacccagttgatcggcgcgagatttaatcgccgcgacaatttgcgacggcg<br>cgtgcagggccagactggaggtggcaacgccaatcagcaacgactgtttgcccgccagttgttgtgccacgcg<br>gttgggaatgtaattcagctccgccatcgccgcttccactttttcccgcgttttcgcagaaacgtggctggcctggt<br>tcaccacgcgggaaacggtctgataagagacaccggcatactctgcgacatcgtataacgttactggtttcacatt<br>caccaccctgaattgactctcttccgggcgctatcatgccataccgcgaaaggttttgcgccattcgatggtgtcc<br>gggatctcgacgctctcccttatgcgactcctgcattaggaagcagcccagtagtaggttgaggccgttgagcac<br>cgccgccgcaaggaatggtgcatgcaaggagatggcgcccaacagtcccccggccacggggcctgccacc<br>atacccacgccgaaacaagcgctcatgagcccgaagtggcgagcccgatcttccccatcggtgatgtcggcga<br>tataggcgccagcaaccgcacctgtggcgccggtgatgccggccacgatgcgtccggcgtagaggatcgag<br>atctcgatcccgcgaaattaatacgactcactataggggaattgtgagcggataacaattcccctctagaaataatt<br>ttgtttaactttaagaaggagatataccatgggcagcagccatcatcatcatcatcacagcagcggcctggtgcc<br>gcgcggcagc |

**Table S3:** Primers used in this study. Primers #1-10 are to clone 12 constructs including four *rppA* coding sequence (native, *E. coli* codon optimized, HC and HR codon harmonized) into three plasmid backbones of pBbE2k, pBbE7k, and pBbE8k. Primers #11-14 are to clone 4 construct including four *rppA* coding sequence (native, *E. coli* codon optimized, HC and HR codon harmonized) into one plasmid backbones of pBbEJ23101k. Primers #15-18 are to clone *E. coli* codon optimized *rppA* plasmid backbones of pET28b.

| # | Primer description | Sequence |
|---|---|---|
| 1 | Native *rppA* Forward | CAAAAGATCTTTTAAGAAGGAGATATACATATGGCGACCCTGTG CCGACC |
| 2 | Native *rppA* Reverse | CCTGGAGATCCTTACTCGAGTTTGGATCCTTATTTTTCGAA CTGCGGGTGGCTCCAGCCGGACAGCGCAACGCCGGCG |
| 3 | Optimized *rppA* Forward | CAAAAGATCTTTTAAGAAGGAGATATACATATGGCTACTC TGTGTCGCCCTGC |
| 4 | Optimized *rppA* Reverse | CCTGGAGATCCTTACTCGAGTTTGGATCCTTATTTTTCGAA CTGCGGGTGGCTCCATCCACTCAGAGCAACTCCGGCGGT |
| 5 | HC *rppA* Forward | CAAAAGATCTTTTAAGAAGGAGATATACATATGGCCACCC TGTGCAGACCG |
| 6 | HC *rppA* Reverse | CCTGGAGATCCTTACTCGAGTTTGGATCCCTATTTTTCGAA CTGCGGGTGGCTCCAGCCGCTCAGGGCTACGCCCGCCG |
| 7 | HR *rppA* Forward | CAAAAGATCTTTTAAGAAGGAGATATACATATGGCAACTC TCTGCCGGCCT |
| 8 | HR *rppA* Reverse | CCTGGAGATCCTTACTCGAGTTTGGATCCTTATTTTTCGAA CTGCGGGTGGCTCCAACCAGAGAGTGCGACACCAG |
| 9 | pBb backbone Forward | GGATCCAAACTCGAGTAAGGATCTCC |

| | | |
|---|---|---|
| 10 | pBbJ backbone Reverse | GCCATATGTATATCTCCTTCTTAAAAGATCTTTTGAATTC |
| 11 | *rppA* Forward | CGTCTTTACAGCTAGCTCAGTCCTAGGTATTATGCTAGCAAGAATTCAAAAGATCTTTTAAGAA |
| 12 | *rppA* Reverse | GCCTGGAGATCCTTACTCGAGTTTGGATCCTTATTTTTCGAACTGCGGGTGGC |
| 13 | pBbJ Forward | GGATCCAAACTCGAGTAAGGATCTCCAGGC |
| 14 | pBb Reverse | GCATAATACCTAGGACTGAGCTAGCTGTAAAGACGTCGGAATTGCCAGCTGGG |
| 15 | Optimized *rppA* Forward | CACACCAGGTCTCACAGCATGGCTACTCTGTGTCGCCCTGC |
| 16 | Optimized *rppA* Reverse | CACACCAGGTCTCATGGTGTTATCCACTCAGAGCAACTCCGGCGG |
| 17 | pET28b Forward | CACACCAGGTCTCAACCACCACTGAGATCCGGCTGCTAAC |
| 18 | pET28b Reverse | CACACCAGGTCTCAGCTGCCGCGCGGCACCAG |

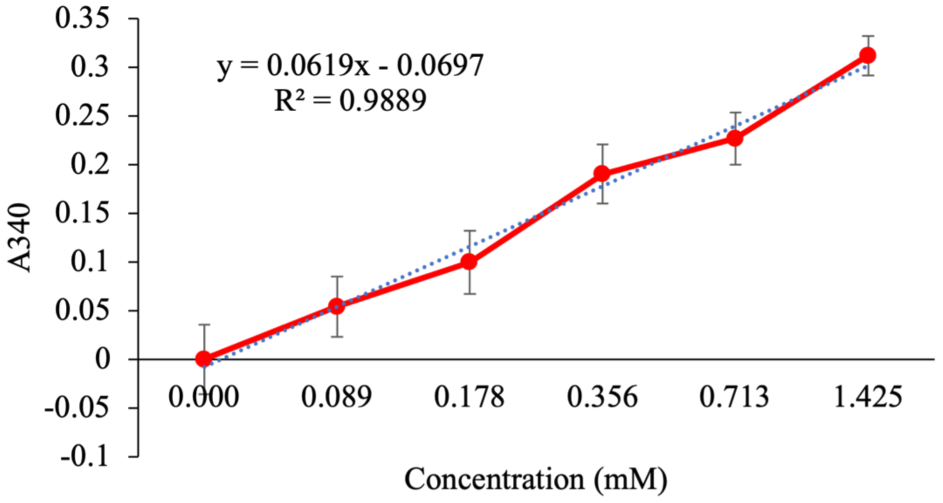


**Figure S1.** Standard curve of flaviolin generated by spiking increasing concentrations of purified flaviolin into BL21 Star (DE3) lysate CFE mock reactions. Absorbance measurements were taken at 340 nm. For visualization of trends, values were averaged and plotted with error bars representing the standard error of the mean (n=3).

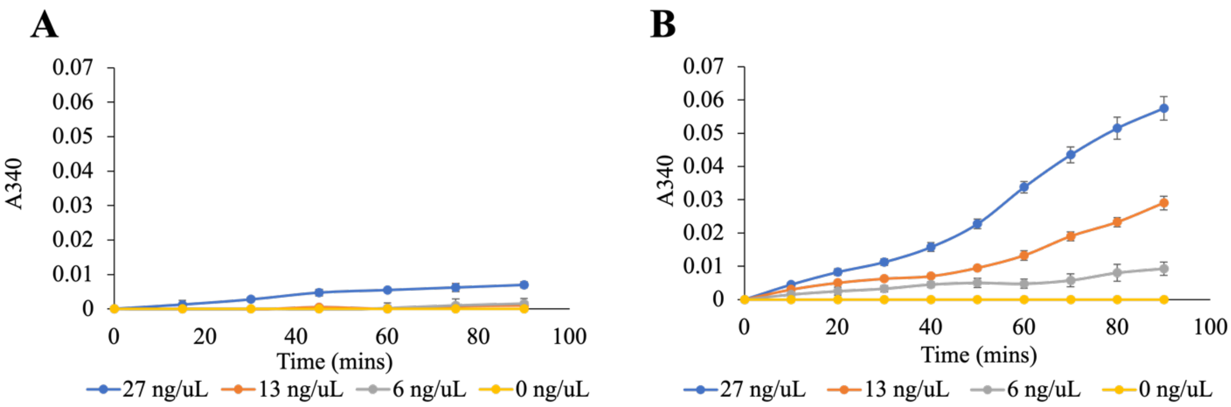


**Figure S2. Initial *E. coli* lysate-based CFE reactions.** Reactions were initiated with increasing concentrations of plasmid DNA at different temperatures to determine the optimal concentration of DNA as well as temperature. (A) CFE reactions at 25˚C. (B) CFE reactions at 30˚C. All reactions were prepared with BL21 Star (DE3) extracts that contained endogenous IPTG and using plasmid DNA of optimized-*rppA* in a pET28b vector. Reactions were run in triplicate and read every 10 mins for 90 mins. Error bars represent the standard error of the mean (*n* = 3).

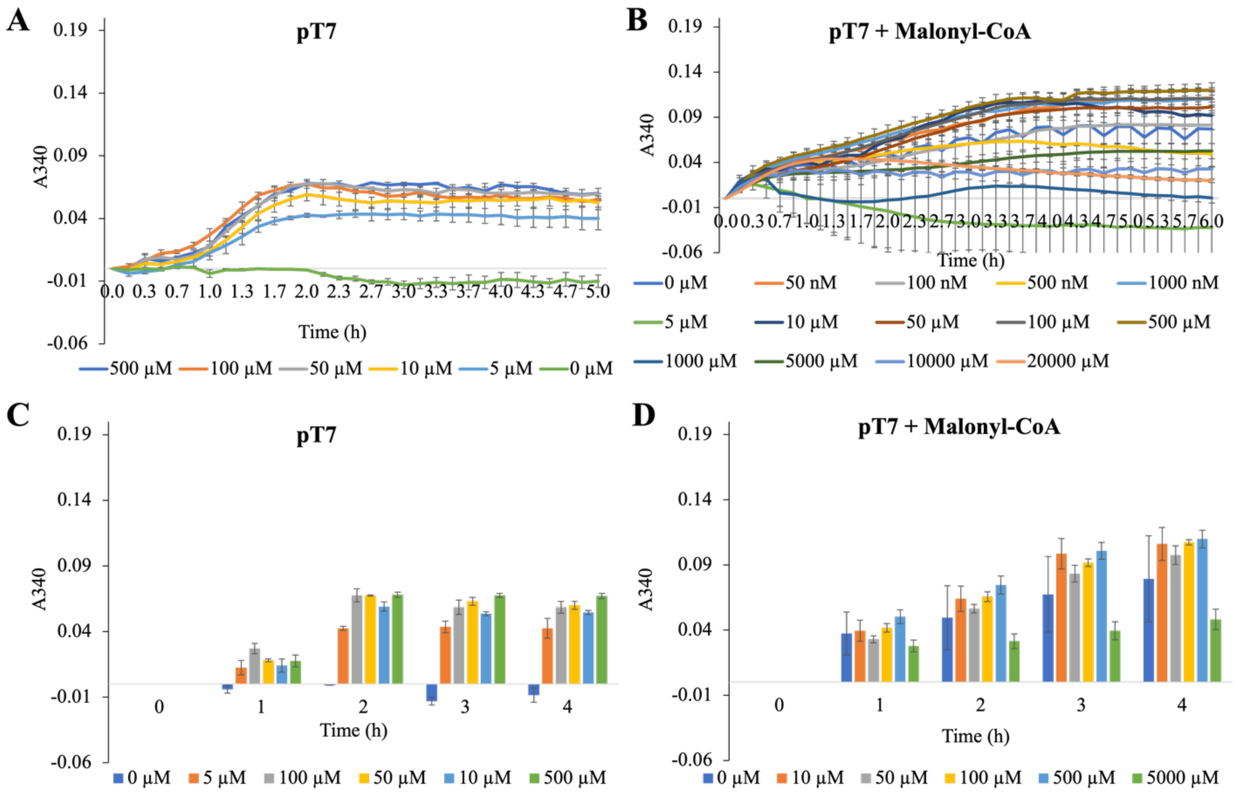


**Figure S3.** Optimization of inducer concentrations in *E. coli* lysate-based CFE reactions for RppA. Reactions were initiated with increasing concentrations of inducers to determine the optimal concentration for the corresponding promoters. All reactions were prepared with BL21 Star(DE3) extracts that lack endogenous IPTG. (A, C) Increasing concentration of IPTG for pT7 promoter using plasmid DNA of optimized-*rppA* driven by pT7 promoter with the addition of 50 ng/µL T7

RNA polymerase. (B, D) Reactions were initiated with increasing concentrations of the extender unit malonyl-CoA concentration to determine optimal concentration using plasmid DNA of optimized-*rppA* driven by pT7, 50 ng/µL T7 RNA polymerase, and 500 µM IPTG. Reactions were run in triplicate and read every 10 mins for 6 hr. Error bars represent the standard error of the mean ($n$ = 3). (C, D) Bar graph of (A, B) respectively.

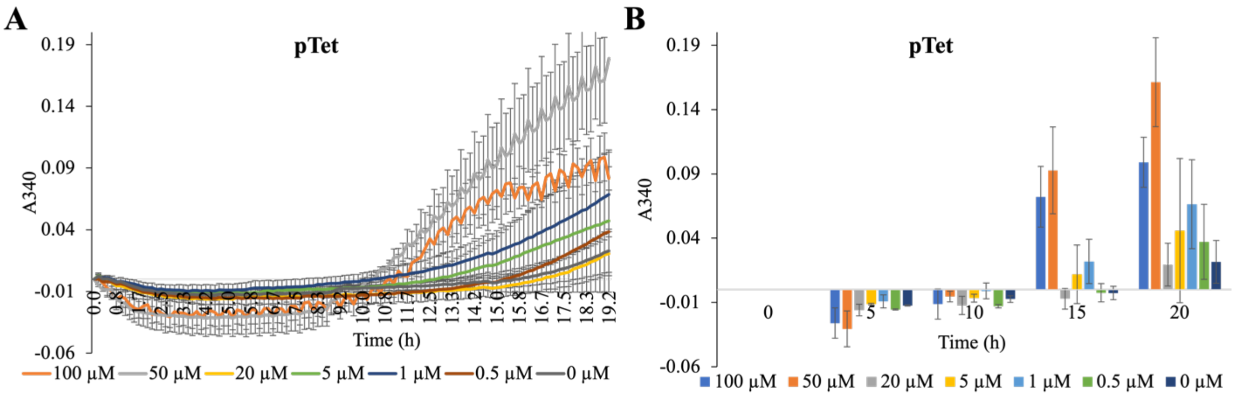


**Figure S4.** Optimization of anhydrotetracycline concentrations inducer for pTet promoter in *E. coli* lysate-based CFE reactions for RppA. Reactions were initiated with increasing concentrations of anhydrotetracycline to determine the optimal concentration for pTet promoter. All reactions were prepared with BL21 Star(DE3) extracts that lack endogenous IPTG, plasmid DNA of optimized-*rppA* driven by pTet promoter, and 500 µM malonyl-CoA. Reactions were run in triplicate and read every 10 mins for 20 hr. Error bars represent the standard error of the mean ($n$ = 3). (B) Bar graph of (A).

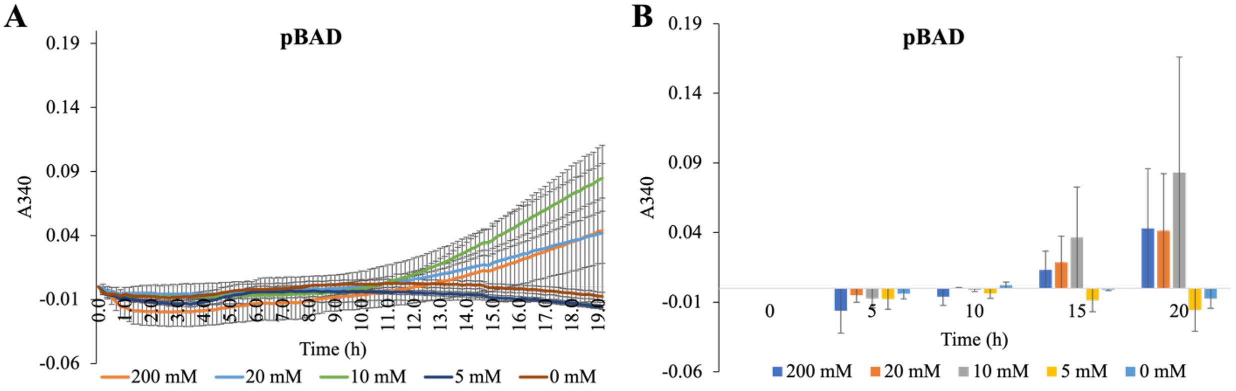


**Figure S5.** Optimization of L-arabinose concentrations inducer for pBAD promoter in *E. coli* lysate-based CFE reactions for RppA. Reactions were initiated with increasing concentrations of L-arabinose to determine the optimal concentration for pBAD promoter. All reactions were prepared with BL21 Star(DE3) extracts that lack endogenous IPTG, plasmid DNA of optimized-*rppA* driven by pBAD promoter, and 500 µM malonyl-CoA. Reactions were run in triplicate and read every 10 mins for 20 hr. Error bars represent the standard error of the mean ($n$ = 3). (B) Bar graph of (A).

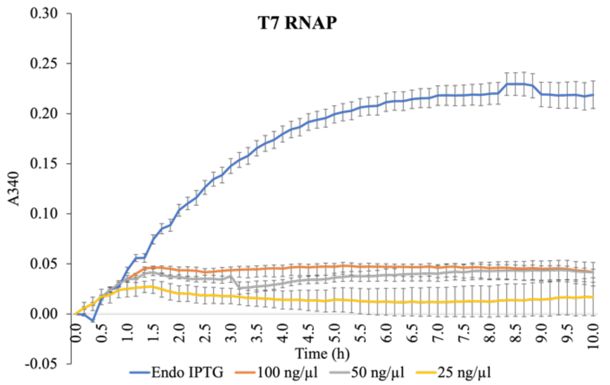


**Figure S6.** pT7 constructs using optimized-*rppA* coding sequence for 1) lysate containing endogenous IPTG, 2) lysate does not contain endogenous IPTG, 500 µM IPTG, with increasing concentration of T7 RNA polymerase. Reactions were run in 6 replicates and read every 10 mins for 10 hr. Error bars represent the standard error of the mean (*n* = 6).

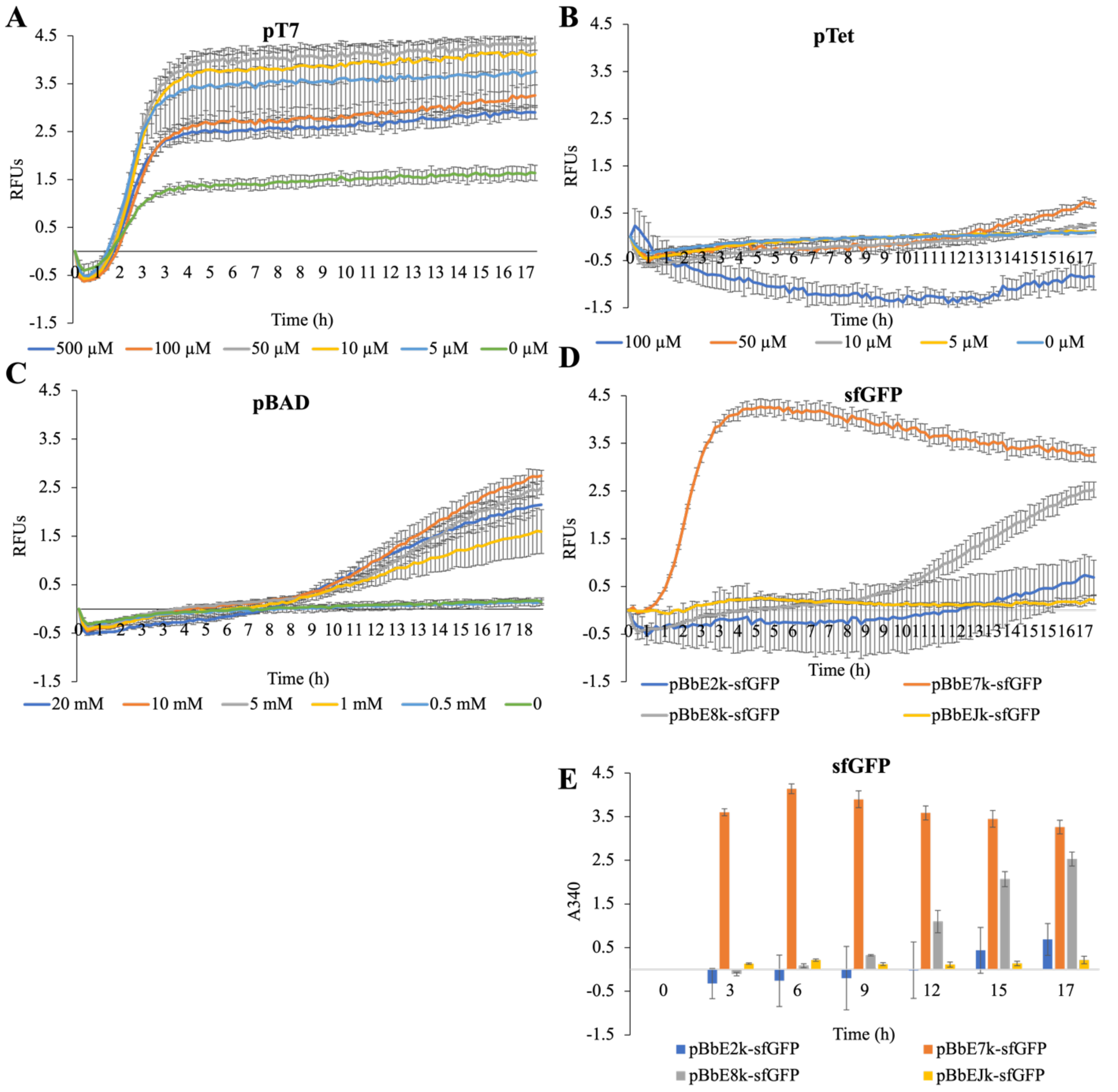


**Figure S7.** Optimization of inducer concentrations in *E. coli* lysate-based CFE reactions for sfGFP. Reactions were initiated with increasing concentrations of inducers to determine the optimal concentration for the corresponding promoters. All reactions were prepared with BL21 Star(DE3) extracts that do not contain endogenous IPTG. (A) Increasing concentration of IPTG for pT7 promoter using plasmid DNA of sfGFP driven by pT7 with the addition of 50 ng/µL T7 RNA polymerase. (B) Increasing concentration of anhydrotetracycline for pTet promoter using plasmid DNA of sfGFP driven by pTet promoter. (C) Increasing concentration of L-arabinose for pBAD promoter using plasmid DNA of sfGFP driven by pBAD promoter. (D) CFE reaction of sfGFP in BL21 Star(DE3) extracts that lack endogenous IPTG. pT7 construct supplemented with 50 ng/µL T7 RNA polymerase and 500 µM IPTG. pTet construct supplemented with 50 µM anhydrotetracycline. pBAD construct supplemented with 10 mM L-arabinose. There is a delay in the expression of pTet and pBAD promoters, and both promoters indicate less expression than the pT7 promoter constructs. Reactions were run in triplicate and read every 10 mins for 20 hr. Error bars represent the standard error of the mean (*n* = 3). (E) Bar graph of (D).

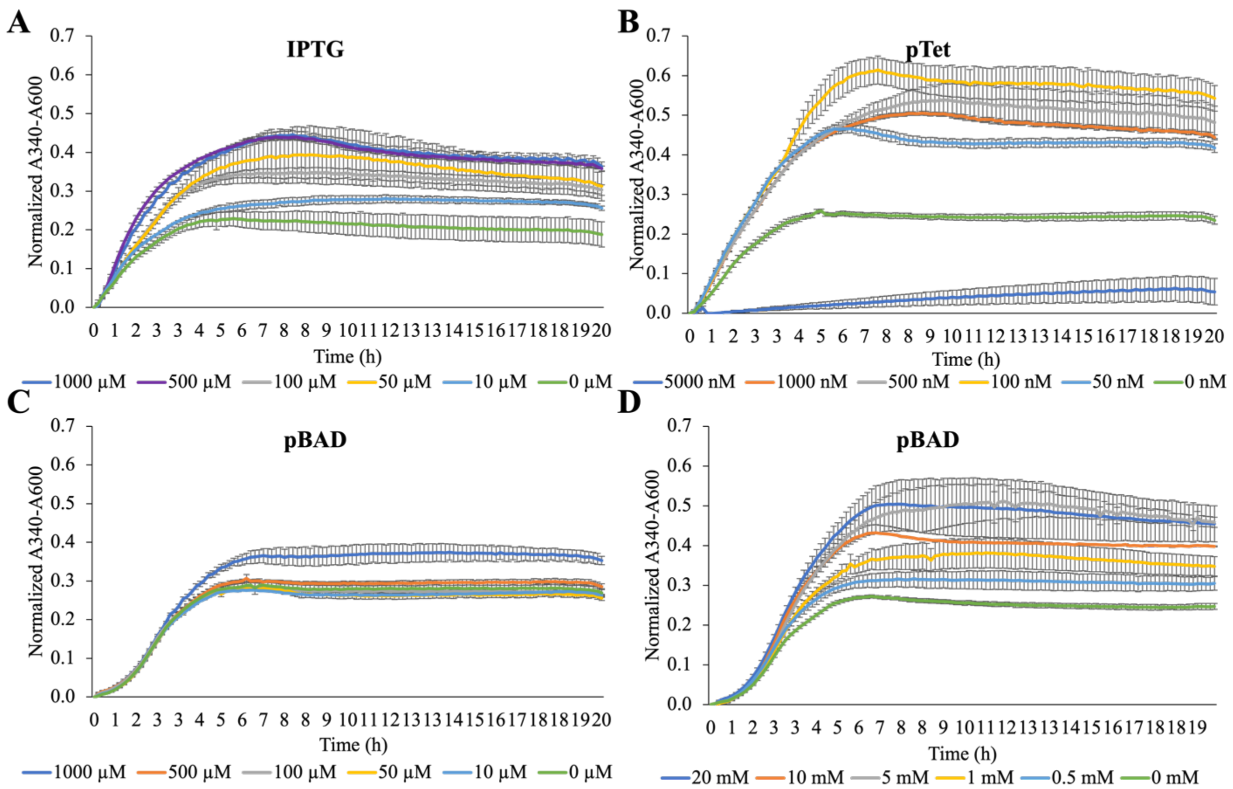


**Figure S8.** Optimization of inducer concentrations in *E. coli in vivo* reactions. Reactions were initiated with increasing concentrations of inducers to determine the optimal concentration for the corresponding promoters. All constructs were expressed in BL21 Star(DE3). (A) Reactions were initiated with increasing concentrations of IPTG using cells harboring plasmid DNA of HG-*rppA* driven by pT7. (B) Increasing concentration of tetracycline for pTet promoter using cells harboring plasmid DNA of HG-rppA driven by pTet. (C, D) Increasing concentration of L-arabinose for pBAD promoter using plasmid DNA of optimized-*rppA* driven by pBAD. Reactions were run in triplicate and read every 10 mins for 20 hr. Error bars represent the standard error of the mean (*n* = 3).
